# Rating Pome Fruit Quality Traits Using Deep Learning and Image Processing

**DOI:** 10.1101/2024.04.03.588000

**Authors:** Nhan H. Nguyen, Joseph Michaud, Rene Mogollon, Huiting Zhang, Heidi Hargarten, Rachel Leisso, Carolina A. Torres, Loren Honaas, Stephen Ficklin

## Abstract

Quality assessment of pome fruits (*i.e.* apples and pears) is used not only crucial for determining the optimal harvest time, but also the progression of fruit-quality attributes during storage. Therefore, it is typical to repeatedly evaluate fruits during the course of a postharvest experiment. This evaluation often includes careful visual assessments of fruit for apparent defects and physiological symptoms. A general best practice for quality assessment is to rate fruit using the same individual rater or group of individuals raters to reduce bias. However, such consistency across labs, facilities, and experiments is often not feasible or attainable. Moreover, while these visual assessments are critical empirical data, they are often coarse-grained and lack consistent objective criteria. Granny, is a tool designed for rating fruit using machine-learning and image-processing to address rater bias and improve resolution. Additionally, Granny supports backwards compatibility by providing ratings compatible with long-established standards and references, promoting research program continuity. Current Granny ratings include starch content assessment, rating levels of peel defects, and peel color analyses. Integrative analyses enhanced by Granny’s improved resolution and reduced bias, such as linking fruit outcomes to global scale-omics data, environmental changes, and other quantitative fruit quality metrics like soluble solids content and flesh firmness, will further enrich our understanding of fruit quality dynamics. Lastly, Granny is open-source and freely available.

## 1 Introduction

The US apple market is worth an estimated $23 billion (USApple, n.d). A large majority (67%) of the ∼11.1 billion pounds of apples produced in the US are intended for the fresh fruit market, yet it is typical for 25% or more to be culled at packout because fruit failed to meet strict thresholds for fruit quality, including cosmetic defects (Gallardo & Pedroso-Galinato, 2020, 2023). Around 80% of pome fruit destined for the US fresh fruit is produced in the Pacific Northwest region of the US (Northwest Horticultural Council, 2023). In order to meet year-round domestic demand for fresh pome fruit, much of that crop is stored, often for several months. During storage, apples and European pears can display a wide range of symptoms that reflect a diversity of physiological disorders and other losses in quality. Therefore, a principle aim of postharvest research is to understand how pome fruits lose quality throughout the supply chain and develop strategies to mitigate these losses. This typically involves repeated evaluation of both visual and physiological characteristics related to fruit quality during the course of experiments. The former is typically performed by skilled technicians who classify fruit by well-established binning schema based on symptom severity. In order to reduce rater bias, the standard best practice is to have the same individual or group of individuals rate fruit throughout an experiment (L. A. Honaas et al., 2016). However, this is not always practical or feasible due to logistical and personnel challenges in a laboratory setting, such as small group sizes within individual research programs, annual employee turnover, variability across locations, *etc*. The eventuality that comparisons of ratings performed by different individuals across years and/or experiments is high, and the inevitable rater bias across programs and years can confound downstream analyses. Moreover, an active area of research in pome fruits is to create machine learning models for fruit postharvest quality traits prediction using as input global scale-omics data and accurate physiological ratings (Washington Tree Fruit Research Commission, n.d). Thus, having access to higher resolution physiological ratings will allow for more nuanced predictive models.

Computer vision has been widely adopted for feature detection from imagery and offers an opportunity to address rater bias in postharvest assessment. Computer vision is already used in the agriculture sector and specifically the pome fruit industry. Notably, applications include the development of image based fruit sorting machines (Rehkugler & Throop, 1986; Wen & Tao, 1999), hyperspectral imaging for fruit quality (Çetin et al., 2022; Nicolaï et al., 2006), robotic harvesting equipment (Bu et al., 2022; Davidson et al., 2016; Hua et al., 2023), and more (Lorente et al., 2013; Pu et al., 2015; Zhou et al., 2023). Importantly, early detection of issues that affect fruit quality is critical for management strategies and minimizing food waste throughout the pre- and postharvest supply chain. Image based analytics have evolved rapidly with recent advancements in Machine Learning (ML) and Deep Learning (DL), and their utilization has gained momentum for image processing in apple fruit analysis (Naranjo-Torres et al., 2020), with specific focus on early disorder detection (Buyukarikan & Ulker, 2022; Mogollon et al., 2020), fruit grading (Bhatt & Pant, 2015; Li et al., 2021; Yang et al., 2022), and in-field yield prediction (Cheng et al., 2017; Datt & Kukreja, 2024). Despite these advances, many fruit quality assessments (such as starch level, peel color, and disorder incidence) are performed manually, in real-world situations and at large scales, likely due to challenges in technology transfer from research and development departments (R&D) to industry, such as workforce knowledge, user friendly interfaces, communication, industry outreach, and a large enough training and testing dataset to make reliable management decisions. There is a need and desire for development of computer vision tools to address rater bias and low resolution for manually rated quality traits (Dhiman et al., 2022).

To address rater bias and improve data resolution, we developed Granny, a modular computer vision software that uses deep learning and image processing to detect fruit in images, and perform rating of disorders or other important fruit-quality traits. We demonstrated the accuracy of Granny by comparing Granny’s fruit quality trait prediction with expert ratings. Finally, Granny is open-source and freely available, and modules from Granny can be adapted beyond pome fruit research, offering potential solutions for various sectors within the agriculture industry.

## 2 Methods

Granny is a computer vision software tool that uses machine learning and image processing to identify individual fruit from photos that contain many (*i.e.* Figure 1A and 1B), then extract individual fruit subimages, and remove the background for later fruit-quality rating (Figure 1C). These ratings include the level of peel disorders such as superficial scald (Figure 1 D1), the peel color (Figure 1 D3 and 1D4), and starch clearing in fruit cross sections (Figure 1 D2).

**Figure 1.**
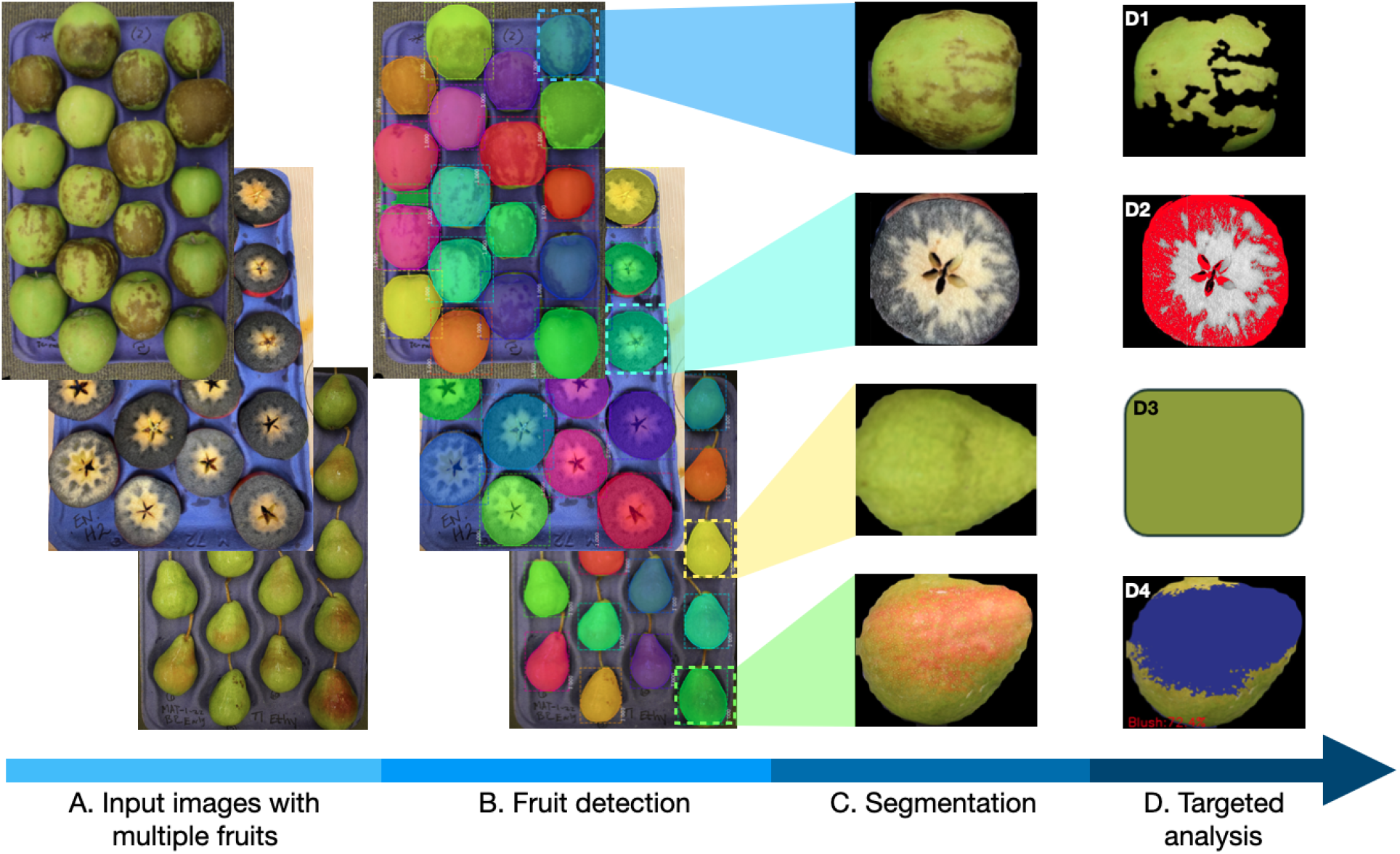
Overview of the automated image analysis workflow. The workflow involves four major steps and the progression is indicated with a blue arrow. **(A)** Examples of input images include trays of apples with superficial scald (top panel), apple cross-sections stained with a iodine solution (middle panel), and pears (bottom panel). **(B)** The fruit detection module identifies individual fruit instances. An enlarged example of the detected instances can be found in Supplemental Figure 1. **(C)** Examples of segmented fruit instances. **(D)** Downstream image analyses include detection and scoring of: D1) superficial scald; D2) starch content; D3) pear background (*i.e.* the shade side) color; and D4) pear blush percentage. For the first three modules, Granny also provides ratings based on user provided references, such as binning scheme for superficial scald, starch rating according to a desired starch pattern index (SPI), and pear color rating according to a pear background color reference card.

### 2.1 Fruit Detection

To identify individuals from images containing multiple fruit instances, a machine learning model, Mask Region-based Convolutional Network (Mask R-CNN), is used (Dollár et al., 2017; He et al., 2020). The Mask R-CNN approach is based on the Feature Pyramid Network (FPN) and a 101-layer Residual Network (ResNet 101) (Lin et al., 2016). Granny employs a publicly available Mask R-CNN model developed by Matterport, Inc (Abdulla, 2017). Initially, the Mask R-CNN detects an object in the image using a previously trained model on a combination of Microsoft Common Objects in Context (COCO) dataset (Lin et al., 2014) and a small balloon dataset provided by Matterport Inc (r v2.1). The pre-trained models were initially used as it provides two key benefits:1) it utilizes the similarity between the ‘round’ shape and colorful filling of fruits and balloons, and 2) pre-trained models save time and computational resources that would otherwise be required for manually training a new model on pome fruits.

While the pre-trained models can detect all the whole fruit instances with high confidence (confidence score >= 0.999), it struggled to identify all the fruit cross-sections on a tray (Supplemental Figure 2). This was likely due to the iodine spills and the shades of the cross-sections on the fiber tray diminishing the contrast between the background and the dark stained fruit tissue, resulting in low-confidence detection and missing instances of cross-sections. To increase the detection confidence for the iodine stained cross-sections, an updated model was trained. First, new image masks of fruit cross-sections were created using the open-source VGG Image Annotator tool (Dutta & Zisserman, 2019). These masks were then used as input to train the updated Mask R-CNN model to extend the initial model. The training was performed for a total of 30 epochs, 100 steps per epoch, a learning rate of 0.001, and the number of proposed classes is 2 (*i.e.* background versus fruit). During the training process, detected fruit with a confidence level below 0.9 were excluded. The trained model for each epoch was saved as a single Hierarchical Data Format (HDF) file, which was later used to improve the detection of fruit cross-sections. This new model extended the initial models (COCO and balloon), and it can be used for detecting both cross-section and whole fruit.

### 2.2 Fruit Image Segmentation

After fruit detection, the input images are segmented into new images of individual fruit utilizing results provided by the Mask R-CNN, including a bounding box, a binary mask, and a detection confidence score (Supplemental Figure 1). The bounding box consists of two pairs of x,y-coordinates (x1, y1, x2, y2). The binary mask is a 2D matrix of ones and zeros, representing the pixel-wise location of fruit tissue in the original image – one mask per fruit. In the mask, a pixel of fruit is represented by a one (1), whereas a non-fruit pixel is represented by a zero (0). The confidence scores are from 0.0 to 1.0, where 0.0 is the least confident in the prediction and 1.0 is the most confident. For confident, positive identification of fruit, Granny’s default segmentation threshold confidence score is >= 0.999. Using the Python OpenCV (Bradski, 2000) and Numpy package (Harris et al., 2020), with the Mask R-CNN results, individual images of fruit are extracted. Granny provides options for controlling the number of extracted fruits. The user can manually specify this number or use the default value (18) to extract the instances with the highest confidence scores. Images of individual fruit are stored in a user specified directory for downstream analyses. The segmented sub-images are named after the original image, with a numeric appendix. For instance, the first sub-images from image A will be named A-1 and the next sub-image will be named A-2.

### 2.3 Apple Superficial Scald Rating

Superficial scald, recognized as brown, necrotic fruit peel tissue, is rated by Granny using Python OpenCV and Numpy. A workflow summary is available as Supplemental Figure 3. Each apple image file (after segmentation) in a given directory is imported into an RGB (red, green, blue) array, which then undergoes the following steps: 1) conversion from RGB to YCrCb color space to remove residual background pixels, 2) smoothing using a 3-by-3 Gaussian kernel to reduce sharp noise (e.g., discoloration in lenticels or small areas of damage), 3) conversion from RGB to CIELAB (also known as the L*a*b* color space) for thresholding scald, 4) creating a binary matrix with 0’s representing scald areas (for pixels above the threshold) and 1’s representing non-scald areas, 5) and performing another smoothing operation on the binary matrix to account for the use of a hard threshold. The threshold in step 3, calculated on the histogram of the a* channel, is determined by the index of the maximum pixel bin along the pixel values and the pixel range of the histogram, detailed in Supplemental Figure 4. For each image, the superficial scald coverage is calculated as the ratio of superficial scald pixels over the total fruit pixels. Each fruit has two images (one each from rotating the fruit 180 degrees). Granny uses the position of the fruit in the first image to identify the same fruit in the second image. The final rating is the average of the two sides of each apple. Outputs include thresholded image files and a comma separated values (CSV) file containing rating for each fruit. To assess performance, superficial scald ratings from Granny were compared with technician ratings on the same batch of 1553 fruits.

### 2.4 Starch Content Rating

Starch content can be estimated from fruit cross-sections treated with a potassium iodide and iodine solution (Blanpied & Silsby, 1992; Figure. 1A & C). This reagent, also known as the Lugol’s solution, is used for the colorimetric detection of starch in organic compounds. The starch content rating module in Granny takes as input, segmented images of fruit cross-sections treated with potassium iodide solution in a given directory. Currently, starch content ratings are provided using an ImageJ (15.3T) macro which automatically quantifies the iodine-stained areas of fruit cross-sections. A summary workflow is available as Supplemental Figure 5.

To estimate starch content, the macro first reads each image into an RGB format. Next, a color threshold which controls the sensitivity of the macro is specified. Default thresholds for apple and pear are provided. Users can perform calibration on a custom dataset to adjust this threshold as needed. After the color threshold is specified, a line bisecting the apple image must be drawn for ImageJ’s ‘region of interest’ (ROI) to properly identify stained regions (see Supplemental Methods). The macro then initializes an ImageJ’s ‘Analyze Particles’ function to find contours from the binary image, where ROIs equal to or larger than 2,000 pixels^2^ are identified. Depending on the threshold settings, the intensity of staining, and the picture quality, small areas of non-stained regions of the fruit may be included in the ROI, resulting in noisy data. To reduce this noise, identified ROIs smaller than 2,000 pixels^2^ are excluded from the ROI measurement. The module then counts the number of ROIs, calculates the sum ROI area, and divides the sum ROI area by the total cross-section area to calculate percent stained area, thus providing an estimate of starch content, and thereby a proxy for starch clearing. Besides calculating starch content, this module rates starch content using established starch pattern index (SPI) scales including cultivar agnostic charts such as the Cornell Starch Index and the ENZA/T&G Global Starch Index, as well as cultivar specific indices such as those for ‘Jonagold’ and ‘Granny Smith’ (details of the SPIs can be found in Supplemental Figure 6), for backward compatibility. Each image from the chosen SPI was assigned a starch score (0-100%) using Granny. Samples are assigned the SPI category with the closest starch score.

To calibrate the threshold and evaluate the performance of the starch quantifying macro, an assessment was created for participants to rate the starch content of iodine-stained cross-sections from 36 apples of mixed cultivars and 36 ‘Gem’ pears. Participants were asked to rank their experience in rating iodine stained fruit as limited (18 participants), novice (9 participants), or professional (9 participants) before starting the assessment to help determine experience based trends. A 15-minute time limit was advised for completion of the assessment to match timeframes experienced in field starch assessment, with starting and ending times logged for all participants. All images were rated from a scale of zero to ten, where ten represented 100% starch coverage (*i.e.* fully stained). A range of thresholds were tested for apples and pears separately, and those that generated ratings most closely fit the mean of the professional starch rating were selected as the default thresholds. After adjusting to the new thresholds, ratings from all participants were compared to Granny’s new ratings to test for correlation.

### 2.5 Pear Color Rating

Pear background color (*i.e.* the shade side of a pear) is an indicator of pear maturity and is often manually measured using a color card (Figure 2A). Granny offers an automated pear background color assessment module using a four-step approach, summarized in Supplemental Figure 7. The first two steps, 1) sub-image background removal and 2) image smoothing, are identical to that in the superficial scald rating module (detailed in section 2.3). In step 3, the processed images are converted from RGB to CIELAB color space where the overall level of greenness and yellowness for each of the images are calculated as channel a*’s and channel b*’s mean pixel values, respectively (Figure 2C and 2D).

**Figure 2.**
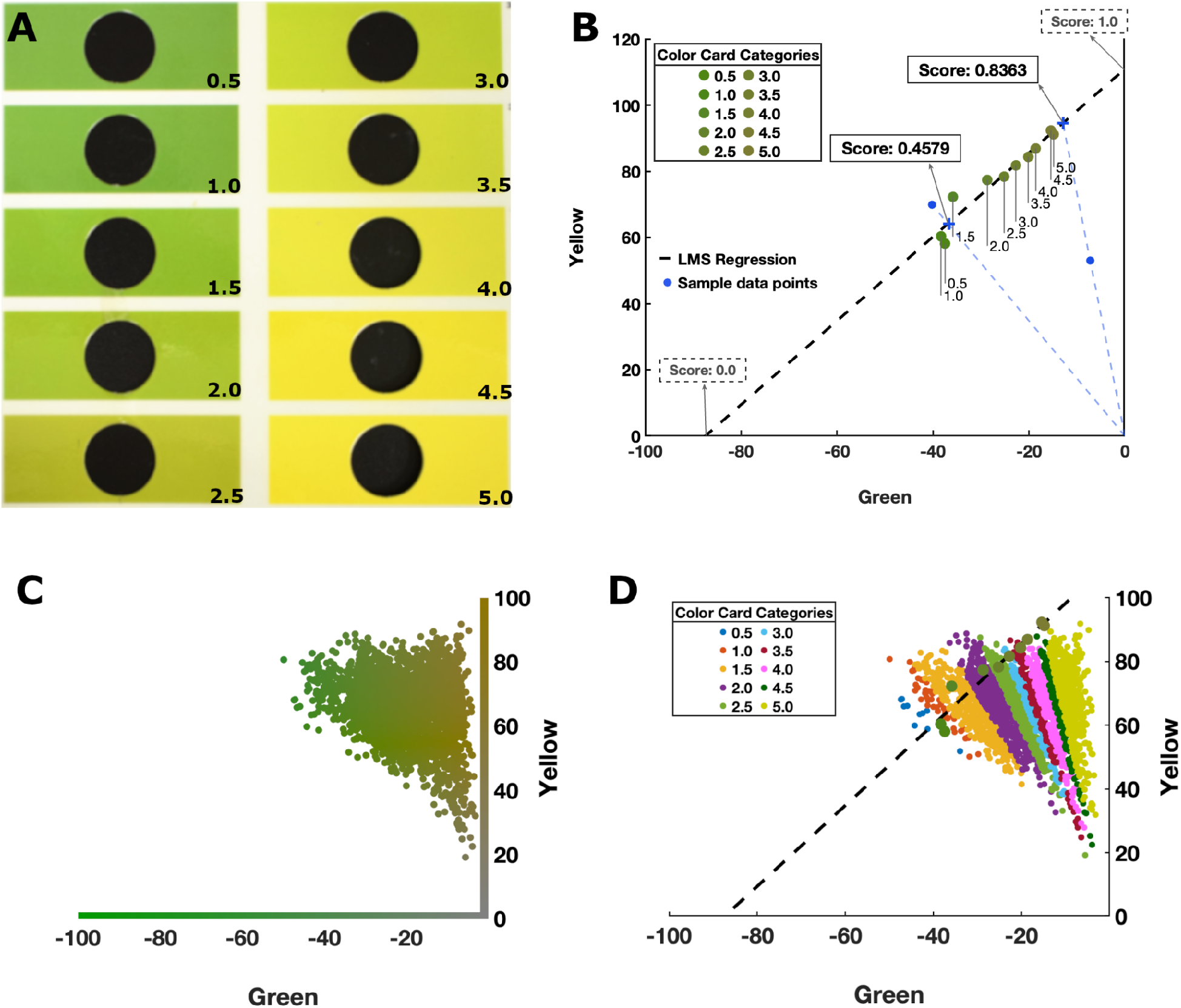
Pear background color rating method and examples. **(A)** A color card widely used in background color rating. It serves as a reference to estimate the level of ripeness of pears based on the ‘greenness’ or ‘yellowness’. **(B)** The 10 colors from the color card are plotted in the CIELAB color space and linear relationship of those colors is demonstrated with the least mean square (LMS) regression line. Color score for a given sample is determined by the value of the intercept (marked with a blue ‘x’) of the LMS line and the line that connects the origin (coordinate: 0.0, 0.0) and the sample point (blue dashed-line). The value of the intercept is determined by its distance from the two ends of the LMS line intercepting either the horizontal axis (score of 0.0) or the vertical axis (score of 1.0). x-axis: a* channel value in the CIELAB color space, representing greenness (green to less green from left to right); y-axis: b* channel value, representing yellowness (yellow to less yellow from top to bottom). **(C)** Calculated overall level of greenness and yellowness of 1,044 samples plotted in the CIELAB color space. Each sample is represented with a dot and is colored with the average color of the sub-image. Axes: same as in (B). **(D)** Same samples (small dots) as the (C) but are colored according to the color bins (big dots) that the samples are assigned into. Axes: same as in (B).

The last step (step 4) calculates a color value based on a standard background color card (Figure 2A). Granny provides two types of color values: 1) a color score ranging from 0.0-1.0, where 0.0 is the greenest color and 1.0 is the yellowest color; 2) a color rating that assigns a pear into one of the 10 color bins from the color card. In the CIELAB space, the 10 background colors from the color card of Figure 2A follow a linear line, fitted by a least mean square (LMS) regression (Figure 2B). Each point along the LMS line is normalized with values between 0.0 and 1.0, where a score of 0.0 corresponds to the location on the LMS line that intersects the horizontal axis, and a score of 1.0 indicates the intercept of the LMS line with the vertical axis. The pear color score is calculated as the value of the point where the line passes through the origin ((0.0, 0.0), lower right corner of Figure 2B) and the pear’s average color of the CIELAB space (blue dashed-line of Figure 2B) intersects the LMS line. Color scores calculated using this method are continuous data.

The color score is then used to approximate the color card rating, which corresponds to the commonly used color card shown in Figure 2A. Granny provides a color card rating (rank of 1 to 10), for backwards compatibility with current practices in the pome fruit industry. However, the color score provides higher resolution for future applications. The color card rating seeks to approximate a similar score as a visual assessment that a trained technician assigns. To do this, color card bins are projected onto the LMS line as described above and each color card bin is assigned a color score. Samples are assigned the color card bins with the closest color score. Figure 2C shows pear colors along the green and yellow axes, and Figure 2D demonstrates how sectors of pears (shown with different colors) map to individual color card ratings. To evaluate the performance of the pear color rating module, color scores generated by Granny were compared to technician ratings from 1,044 pear fruits.

### 2.6 Blush Detection and Quantification

The blush application implemented in Granny takes a 2-step approach to detect and quantify peel blush. First, the ‘cal_blush’ function allows users to define a threshold for quantifying blush color. This is also referred to as the calibration step. This user defined threshold is then used in the ‘blush_percentage’ function to quantify the blush color over segmented images.

In addition to automated image processing steps, the calibration function requires user input for decision-making. Unlike the other modules where a predetermined threshold was used or an automated thresholding method was applied, a manual calibration function was implemented. Various factors, such as production practices and marketing strategies, have significant impact on determining blush thresholds, even for the same batch of fruits. To initiate the calibration, the user first selects a subset of three segmented images from their pool of input images that represent different blush severity: no blush, light, and intense-colored blush (Supplemental Figure 8A). These images are then resized to 500 x 300 pixels each and converted from RGB to CIELAB color space. Next, the three images are concatenated according to blush intensity, then framed in a Python openCV window, where the images are ordered from left to right (no blush to intense blush). This window includes a slider-bar at the bottom of the image, which can be used to manually change the a* channel threshold (min: 0, max: 255). A transparent mask created from each original image is used to over color the pixels where the a* channel detects blush color (Supplemental Figure 8B and 8C). The mask image overlaps the original image and is refreshed automatically as the user manipulates the a* channel with the slide bar to achieve the desired mask overlay. The selected a* channel threshold will be automatically saved upon exit.

After calibrating to the desired a* channel threshold, the ‘blush_percentage’ function is used to quantify blush percentage in a set of segmented images. The ‘blush_percentage’ function reads through each image in a given working directory and converts the images from RGB to the CIELAB color space. For each image, the number of fruit pixels is determined by the number of pixels with a color different from the black background color, and the number of blush pixels is defined by the sum of pixels with a value higher than the a* channel threshold defined by the user during the calibration process. The blush percentage is calculated as the ratio between fruit over blush pixels (number of blush pixels / number of total fruit pixels). The ‘blush_percentage’ function returns an RGB image copy of the segmented image with the blush pixels overlaid in purple and annotated with the percentage of blush on the fruit (Figure 1 D4). A table is generated with information of number of fruit pixels, number of pixels with blush, and blush percentage of each image. Granny blush scores were compared to technician ratings on a set of 175 pear images with different blush coverage. Three expert fruit quality technicians ranked the blush percentage of the original pear images displayed on an HP 24yh monitor (default settings).

## 3 Results

### 3.1 Fruit Detection and Segmentation

The initial fruit detection model trained with the Microsoft Common Objects in Context (COCO) dataset plus a small balloon dataset had a high success rate in detecting fruit instances with high confidence (Table 1). This initial model detected all 10,314 apple and pear instances (100% success rate) and 10,274 of those (99.61%) were detected with strict confidence score threshold (>=0.999). However, the performance of the initial model was unable to detect ∼3% of the tested iodine stained cross-sections and only ∼30% of the detected cross-sections passed the strict threshold. The extended model, which incorporated a custom training dataset of cross-sections, was able to detect 100% of the cross-sections with high confidence. Although both models detected non-target objects as targets, none of them passed the confidence score threshold.

**Table 1.**
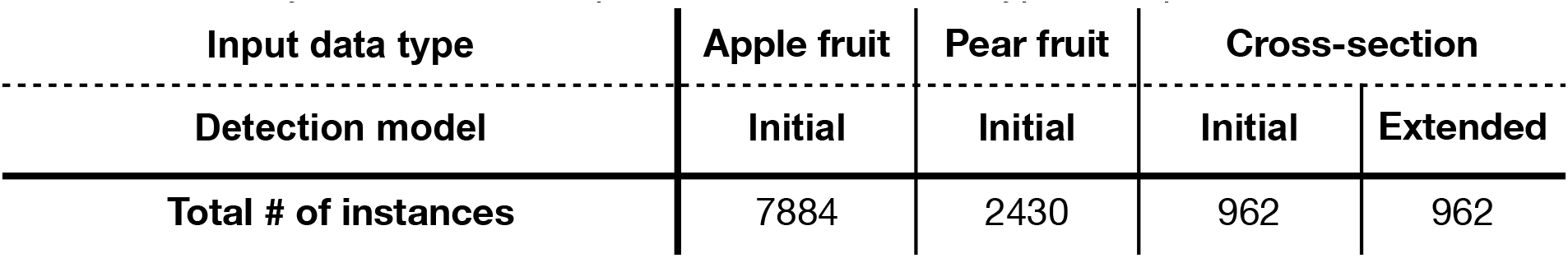
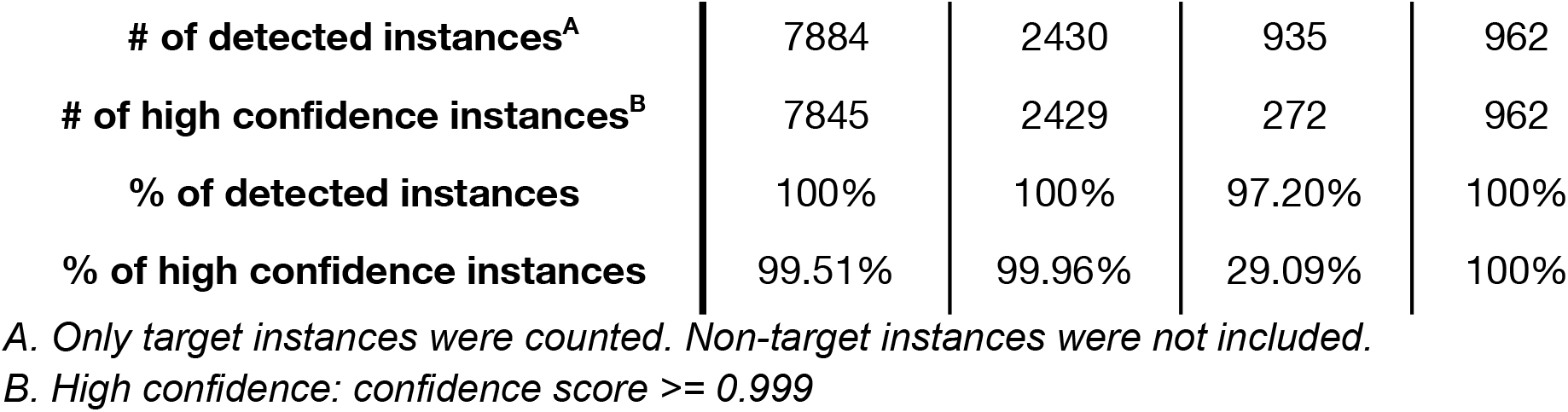
Summary of fruit detection performance with three types of input data and two models.

### 3.2 Superficial Scald Rating

The performance of the superficial scald rating module was evaluated by comparing the predicted scores to a human technician rating (Figure 3). The technician rated fruit into bins (1 to 5), classified by percentage coverage of scalded tissue (0 being no scald and 5 being complete scald), while Granny provided scores ranging from 0.0 (no scald) to 1.0 (complete scald). Generally, scores from Granny were comparable with the technician rating, namely fruits rated with a higher level of superficial scald coverage by technicians are also assigned a higher scald score by Granny. Notably, scald coverage estimations from Granny were lower than human ratings. For example, technicians classified fruits with over 75% scald regions into bin 5, while the same fruits were assigned with a mean score near 61% and upper and lower quartiles near 50% and 70% respectively. Such a trend was generally true across all the bins.

**Figure 3.**
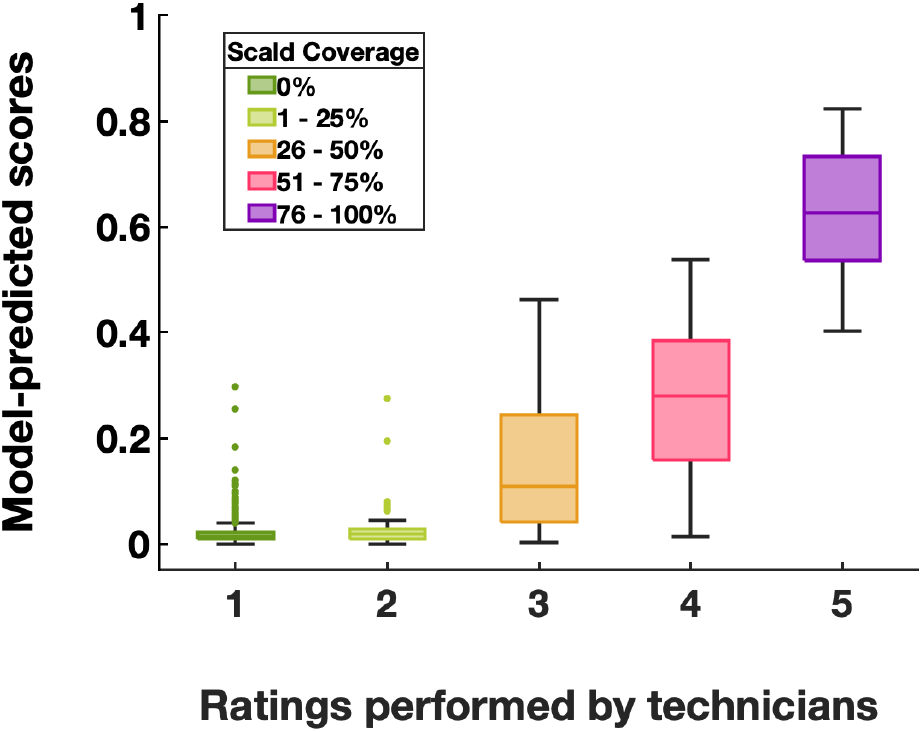
Distribution of the Granny predicted scores (y-axis) for each technician scald coverage bin (x-axis). Fruits were rated into bins (1-5 on the x-axis) by technicians based on the percentage coverage of scalded tissue (1 = 0%, 2= 1-25%, 3=26-50%, 4=51-75%, 5=76-100%). Granny predicted scores (ranging from 0.0-1.0 representing no scald to completely scalded, respectively) for fruits in each technician rated bins were summarized in box plots. A total number of 1,553 ‘Granny Smith’ fruits were assessed.

### 3.3 Starch Content Rating

Starch rating of fruit cross-sections is a commonly used method to estimate pome fruit maturity, especially for apples. The performance of the starch rating module was tested by comparing the percentage of iodine stained cortex tissue estimated by Granny and that by humans. Starch estimations conducted by participants in different experience groups showed high amounts of variability in rated starch content (Supplemental Figure 9). However, the average visual assessment score of all experience groups closely correlated with the starch quantifying module implemented in Granny (Figure 4).

**Figure 4.**
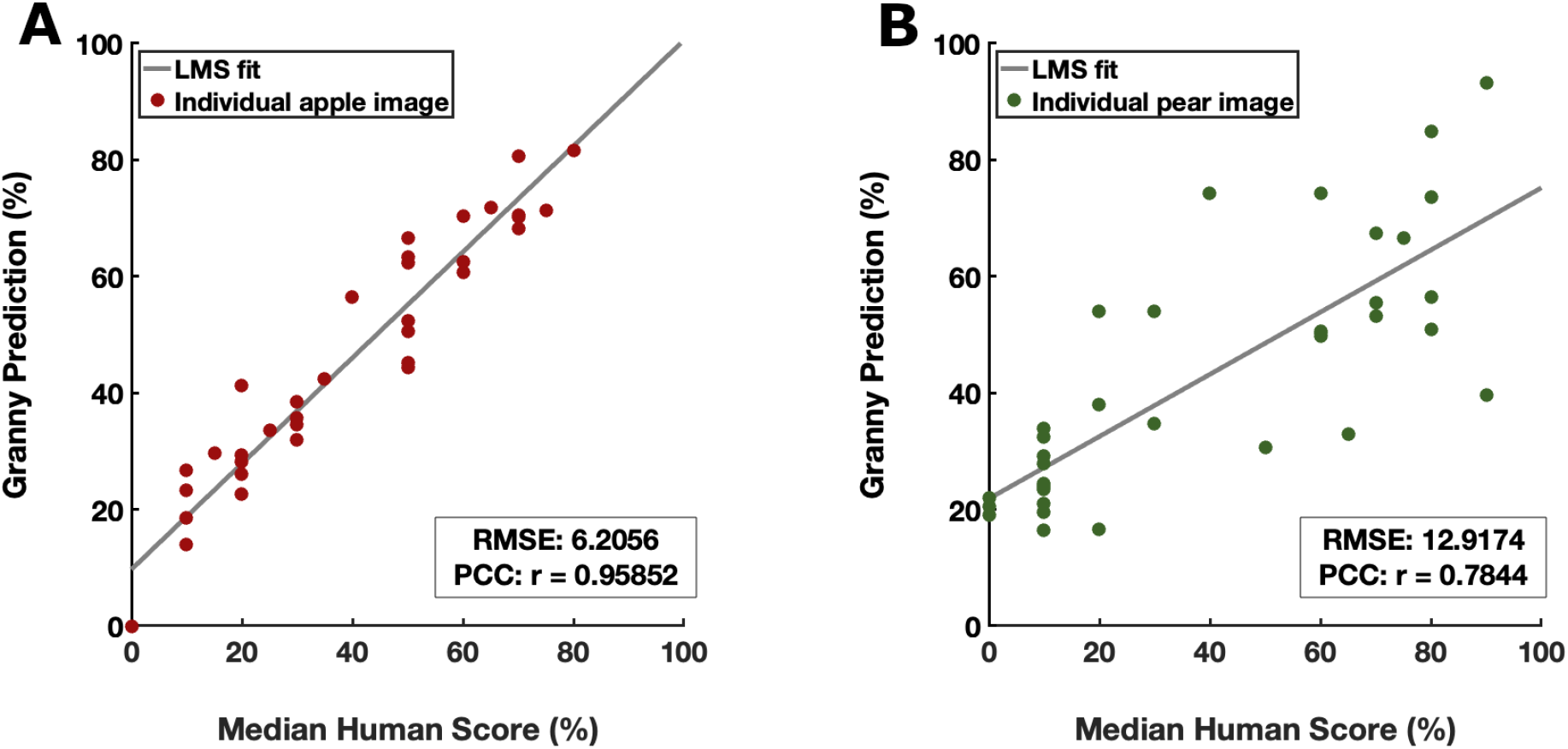
Starch content estimation from Granny and human participants. Comparison of automated starch scores generated by Granny (y-axis) with scores from human participants (x-axis) for apples **(A)** and pears **(B)**. The Least Mean Square (LMS) fit represents a linear relationship between starch scores predicted by Granny and human participants. 36 apples from a variety of cultivars and 36 ‘Gem’ pears were evaluated. RMSE: Root Mean Squared Error. PCC: Pearson Correlation Coefficient.

### 3.4 Pear Color Rating

To determine the accuracy of Granny’s pear color rating module, the automated color scores were compared against expert technician ratings (Figure 5). In summary, the color scores obtained by Granny corresponded to technician’s ratings. Generally, fruits rated higher by a technician also tended to receive a higher score from Granny, both indicating a pear with a deeper yellow color. While Granny’s scores generally align with technician’s ratings, outliers were observed in the first two technician levels (0.5 and 1.0). Examination of the original pear images suggested that the discrepancy was caused by human error as shown in Supplemental Figure 10: the pears classified into the first two bins appear to be more yellow in color than the categories 0.5 and 1.0.

**Figure 5.**
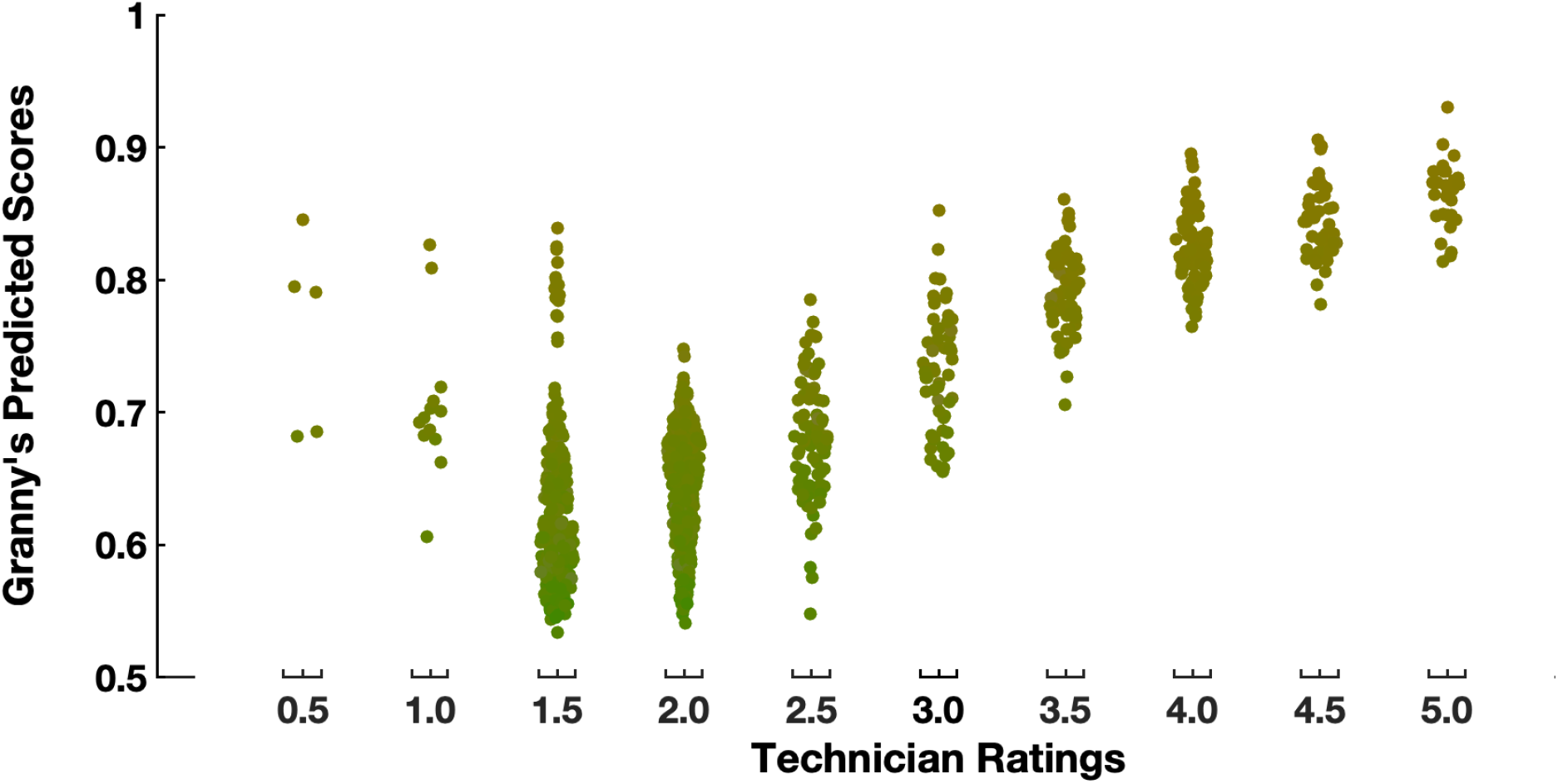
Relationship between the expert technician ratings and the machine-calculated scores. Each dot represents one sample and is colored with the average color of the sub-image estimated by Granny. A total of 1,044 samples are plotted. The x-axis is divided into bins matching the categories from the background color card (Figure 2A). The y-axis is color scores from Granny.

### 3.5 Blush Detection and Quantification

The pear blush detection algorithm allows users to set custom thresholds to distinguish between blush and non-blush regions in pear images, and calculate the proportion of blush regions on each pear (Figure 1D4). Although notable differences are observed in technician ratings on the same fruit image, the average blush percentage estimated by technicians and predicted by Granny had a positive linear correlation (Supplemental Figure 11).

## 4 Discussion

Granny is expected to improve postharvest research by providing automated and reproducible fruit quality ratings. The three major advantages for Granny are first, removal of rater bias, second an increase in the granularity of ratings, which will be important for research studies where higher resolution is necessary, and third, backward compatibility with current standards via mapping of ratings to commonly used standards (*e.g.* starch cards and peel color cards). Currently, Granny provides four rating modules: superficial scald, starch clearing, peel color, and percent blush. Additional comments about each of these modules, including some limitations, are provided in the following sections. In general, we note that because human ratings have bias, we lack “gold standard” ratings by which we can evaluate true performance. Even though expert raters do not provide exact ratings, human ratings and Granny ratings show concordance.

### 4.1 Scald Rating

Superficial scald ratings of ‘Granny Smith’ fruit were quantified after thresholding “non-scald” image pixels using a histogram, which is computationally quick. Because this rating module quantifies the necrotic (*i.e.* brown) portions of green apple fruit peel, it may incorrectly classify other darker regions as superficial scald. This can include other necrotic peel disorders, peel discoloration from mechanical damage, and other parts of the fruit like the pedicle. Until Granny can be improved to distinguish those aforementioned features, it is recommended that users exclude from scald rating, images of apples that have higher incidence of other features. For apple images containing primarily scald, the tool was able to accurately reproduce human ratings of fruit.

### 4.2 Starch Rating

Starch clearing is a widely used index for apple fruit maturity (Blanpied & Silsby, 1992). While the extent of starch clearing is relatable to fruit maturity across apple cultivars generally, there are cultivar-specific characteristics for the rate of clearing, patterns of clearing (*i.e.* across equatorial sections of apple fruit), and the extent of clearing associated with a given fruit maturity level. Many cultivars have dedicated starch index rating schemes that are used to estimate harvest dates (Supplemental Figure 6), but where these dedicated schemas are not available, growers substitute others. A unifying feature of starch iodine indices is that they consist of ∼5-10 discrete categories, but it is not uncommon for users to sort values into sub-categories. This high degree of variability across cultivars and individual usage compounds the issues of rater bias and low starch index granularity. The starch rating module of Granny addresses each of these limitations by quantifying starch clearing in pome fruit cross-sections with a high degree of accuracy compared to human ratings, and greater granularity than human ratings. Further, the software sorts fruit images into existing starch indices and we provide a method to add other starch indices as users desire. This allows backwards compatibility where existing starch indices have been used in the past and cross compatibility where different indices have been used, thus enhancing comparisons across experiments. Digital fruit images can also be reclassified with Granny into a new starch clearing schema as they are developed.

Similar to the superficial scald rating module, other image regions can be misclassified as ROIs of starch clearing (or conversely staining). Yet the software recapitulates human ratings with high accuracy, indicating that these misclassifications are a minor issue for our pilot dataset. Interestingly, human ratings were sometimes consistently high, indicating a minor but systematic bias between humans and Granny (*e.g.* most samples are located above the 1:1 line in Supplemental Figure 9A, C, E, indicating a higher score from human raters). Future work to understand the cause of differences between humans and Granny ratings could inform ways to improve both human ratings and the Granny rating algorithm.

### 4.3 Pear Color

Assessing pear maturity has been a challenge due to the lack of an easy, reliable, and measurable trait that reflects maturity, such as starch clearing in apples. One common measurement taken by scientists and producers that can help estimate pear maturity is the background color, which usually changes from green to yellow as the fruit ripens. Peel color is therefore an important trait that is scored visually in postharvest research. Similar to starch indices where fruit are sorted into a small number of (often commodity or cultivar-specific) color categories, fruit raters use a visual reference card to score individual fruit. Defining the color categories for the card used in this study required an orthogonal classification within the yellow/green colorspace. This revealed narrow and irregularly-spaced categories of peel color that human raters were required to use. We note that it can be challenging to score European pear fruit into these categories by hand due to the narrow color categories (see Fig 2A & B).

Granny can assign fruit by an objective color space value with high granularity and accuracy, potentially offering superior rating performance when compared to hand ratings with the use of a color card. As shown in Figure 2D, pears do not all appear near the line passing through the color card levels. Some appear well below the line. The color rating module will also provide a second metric: the distance of the pear from the line. More work is needed to determine if a higher distance indicates lower quality color ratings. For now, users can treat pear images with a high distance as potential outliers that need further exploration.

As is common with many color processing and matching techniques, variations in lighting conditions and suboptimal image quality can introduce significant fluctuations in the extracted green-yellow values, potentially leading to deviations from technically accurate color ratings. Though this is not a limitation *per se* because human vision where scoring fruit peel color is easily influenced by the lighting conditions.

### 4.4 Blush Detection and Quantification

Classification systems for the presence of blush (red peel coloration) in pome cultivars like ‘Granny Smith’ apples or ‘d’Anjou’ pears are lacking. Therefore, an approach is needed that can incorporate blush into the assessment process alongside other fruit quality data. This is particularly important because blush can significantly impact consumer acceptance. Human visual blush evaluation can be biased. This bias was confirmed with the observation made by three experts on the same fruit where standard deviation was as high as ∼26% in one case (Supplemental Figure 11 E and F). In some cases, the human visual evaluation can be biased based on the blush intensity rather than the total area affected. Granny removes this bias by allowing users to define blush intensity levels through computer vision controlled with a chosen threshold. The current version of Granny provides a rater calibration step, where users provide three images representing extremes. In future versions of Granny we aim to add in an automated threshold detection option similar to the scald rating module.

### 4.6 Limitations and Future Work

Currently, the Granny software is the product of multiple efforts from three different research groups. We determined that joining our efforts into a single software product would be better for management of the software and accessibility to others. The side effect is that the current version of Granny does have a mix of technologies and designs. For example, Granny is written primarily in Python, but the starch rating is provided by an ImageJ macro. Also, the manual thresholding step for the blush module is not consistent with the automated thresholding of the scald module. These differences do result in a non-uniform user experience, however, we are actively working to address these inconsistencies. Future versions of Granny will follow a modular design and will provide a common user experience.

## Availability Statement

Project name: Granny

Project home page: https://github.com/SystemsGenetics/granny Operating systems: Platform independent

Programming language: Python and ImageJ macro

Any restrictions to use by non-academics: GPL v3.0 license

## Acknowledgments

The authors would like to acknowledge Allan Bros., Inc. for providing fruit pictures, USDA-ARS NP-306 Project No. 2094-43000-008-000D, (CT) USDA-NIFA (Hatch project 1014919 (WSU ref. 00011) and Hatch Multistate project NE-2336).

## Author Contributions

SF, LH, RL, and CT conceived the project and acquired funding. NN, JM, RM, HZ, and HH conducted the experiments, gathered data, and performed analyses. All authors contributed to the writing and editing of the manuscript.

---

**Supplemental Figure 1.**
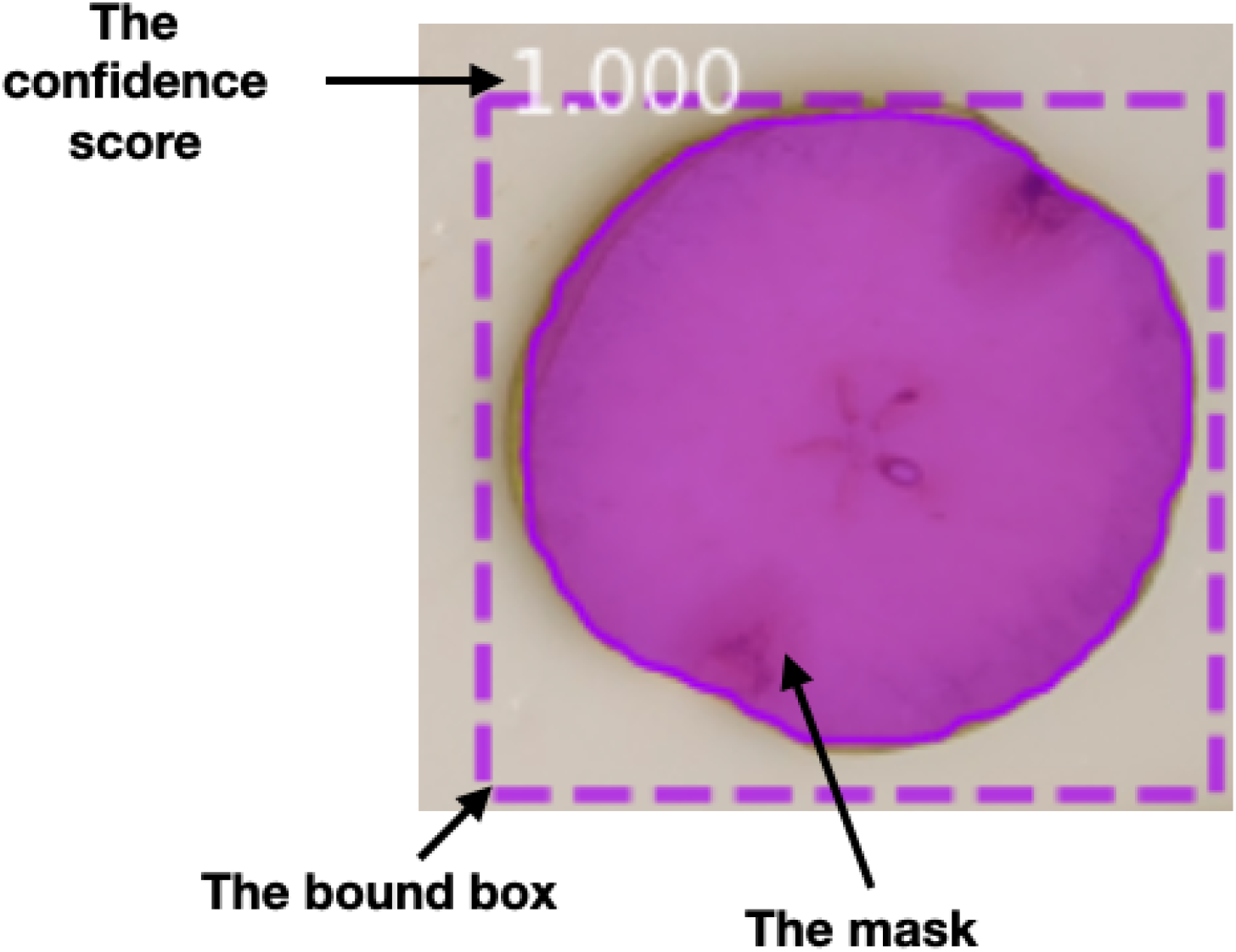
The fruit detection module creates a segmentation mask (colored overlay with solid outline), a bound box (correspondingly colored rectangle with dotted outline), and a confidence score (number shown on the up-left corner of the bound box) for each detected instance. These data will be used for segmentation of the instance.

**Supplemental Figure 2.**
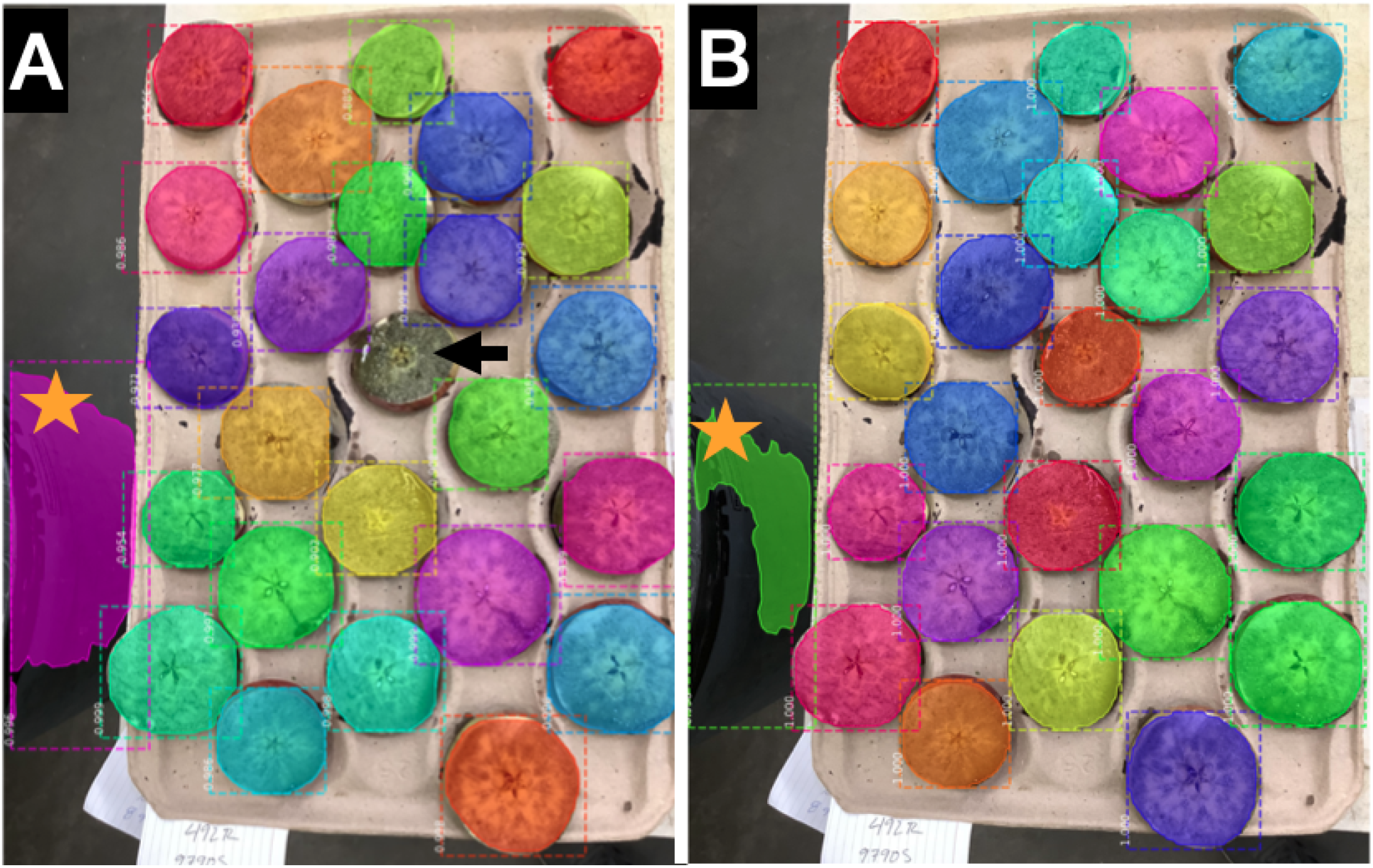
Examples of fruits detected with the initial model **(A)** and the extended model **(B)**. One piece of cross-section was missed in the initial model (indicated with a black arrow in **(A)**), but is detected with high confidence with the extended model. Although the same false positive region (indicated with yellow stars) was detected in both models, the confidence score is lower in the extended model (0.990) than that in the initial model (0.996)

**Supplemental Figure 3.**
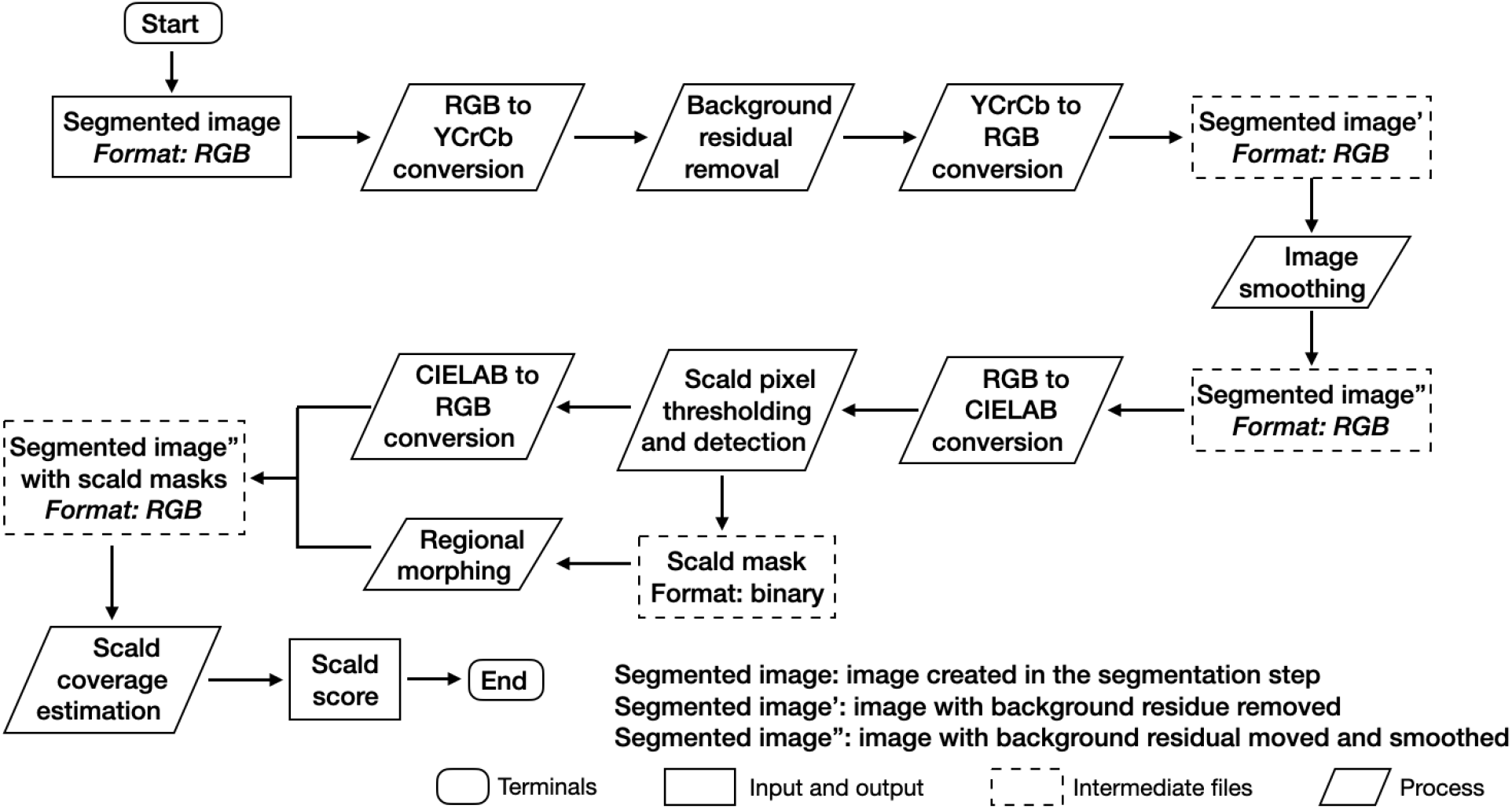
Superficial scald rating workflow

**Supplemental Figure 4.**
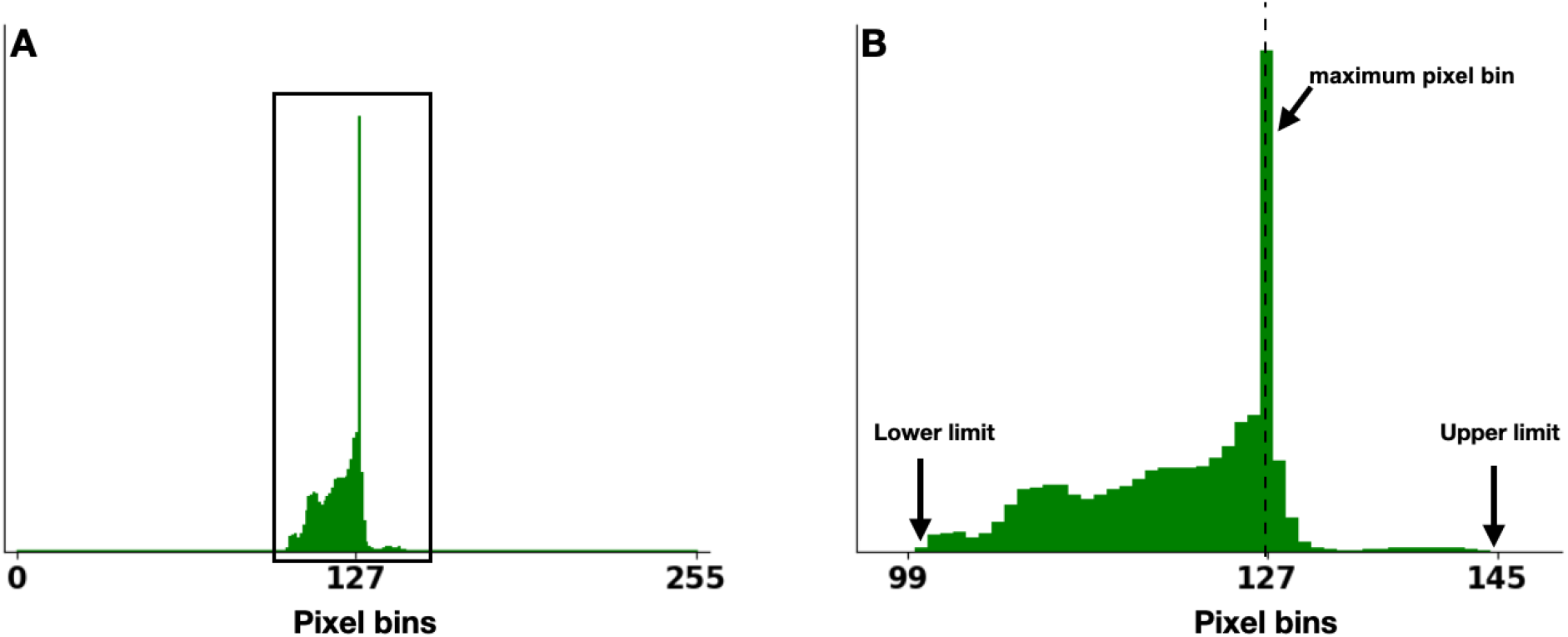
Histogram along the a* channel in the CIELAB space of an apple image for demonstrating the scald thresholding method. (A) shows the 0-255 range of the a* channel. Pixel bins increment by 1 from 0 to a maximum of 255. The region in the black box in (A) is expanded as (B). The lower limit is defined as the pixel bin with the lowest pixel values in the image, whereas the upper limit is the pixel bin with the highest pixel value. The maximum pixel bin is defined as the bin with the most pixels. X-axis shows the range of pixel bins, y-axis is the relative abundance of pixels in each bin. The definitions of histogram, range, and pixel bin are adopted from: https://docs.opencv.org/3.4/d8/dbc/tutorial_histogram_calculation.html Below shows the methods for threshold determination: Apple pixel range = upper limit - lower limit Threshold = maximum histogram pixel bin - 1/3 apple pixel range For the example shown in here, the threshold is calculated as below: Threshold = 127 - 1/3 (145-99) = 111.67

**Supplemental Figure 5.**
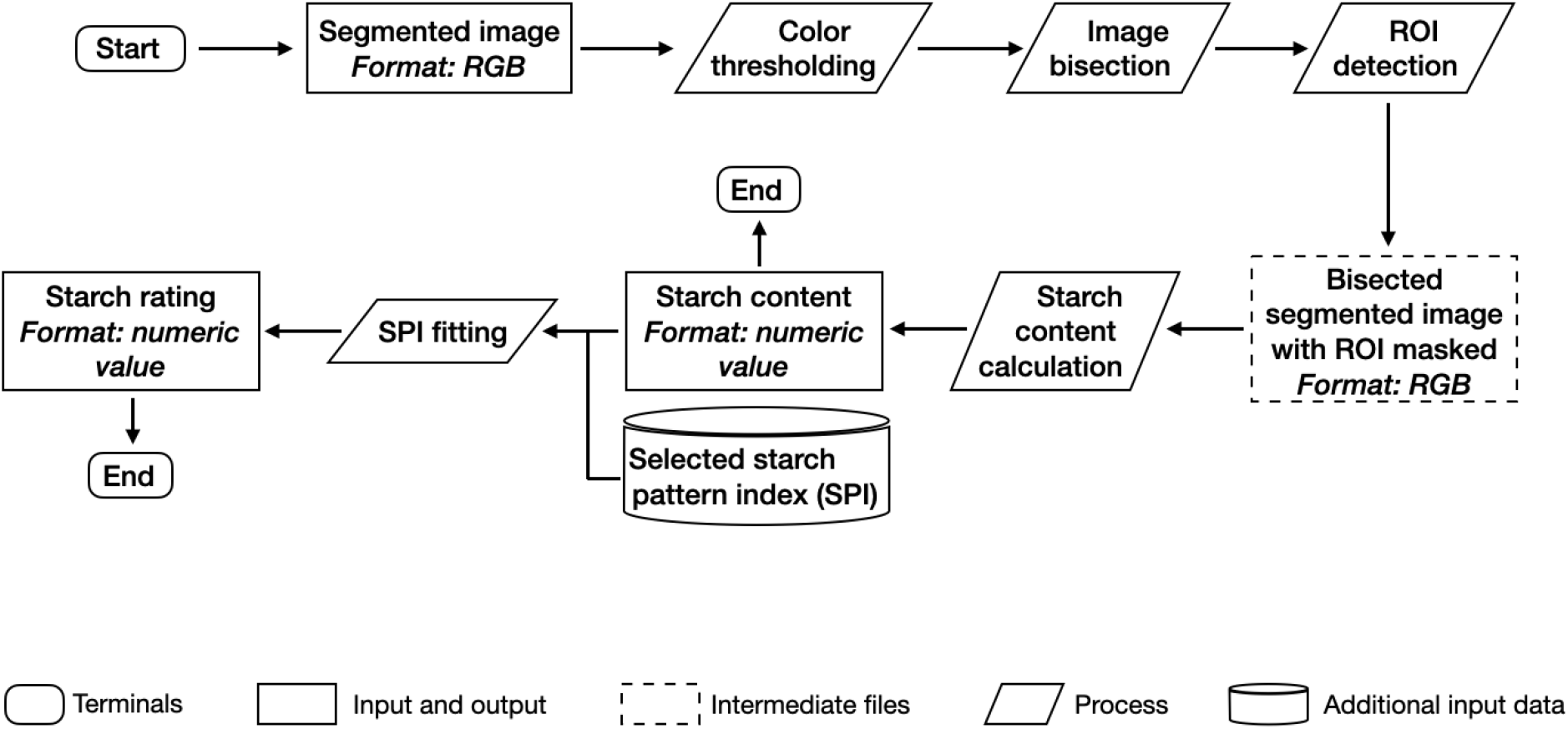
Starch rating workflow

**Supplemental Figure 6.**
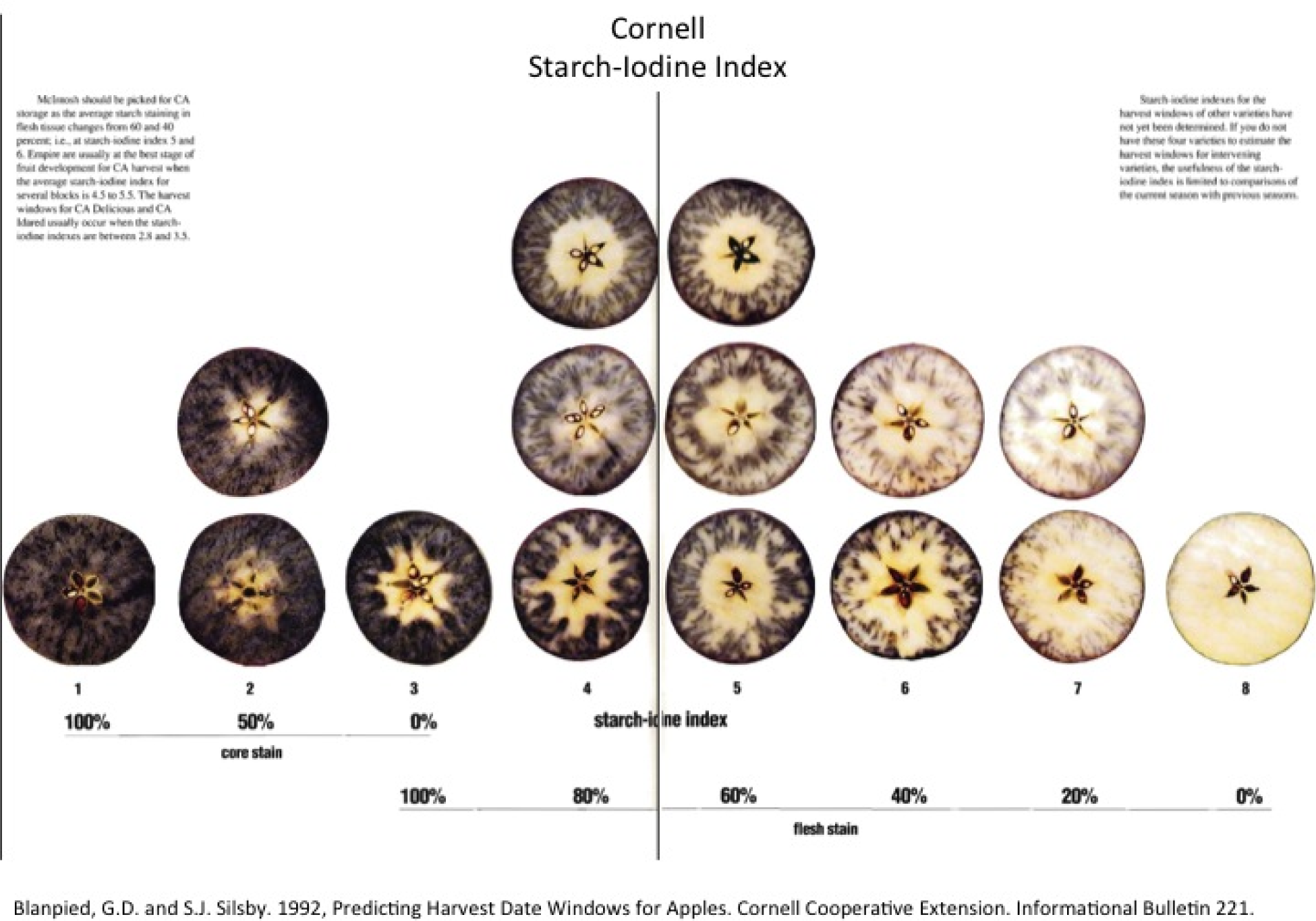
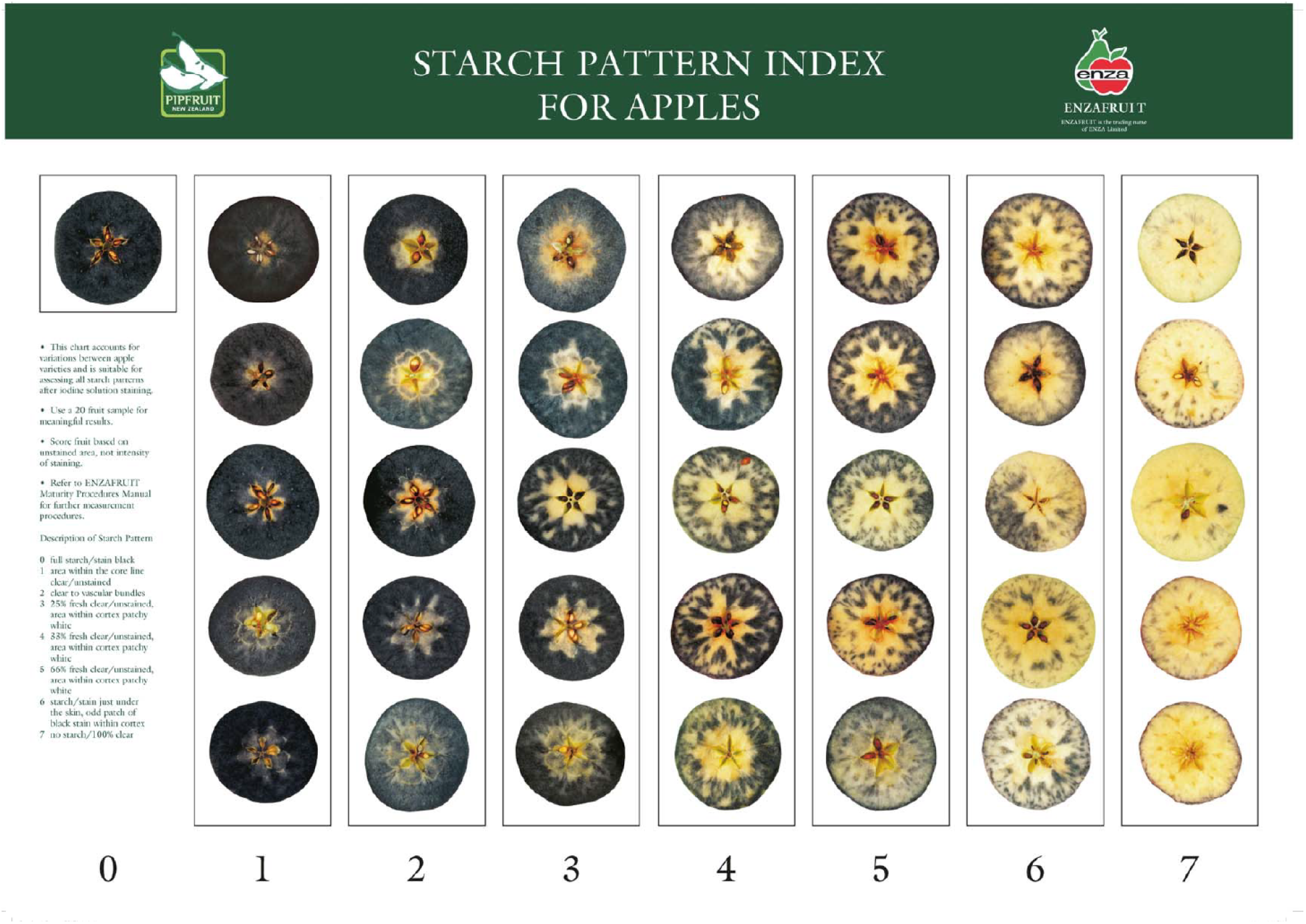
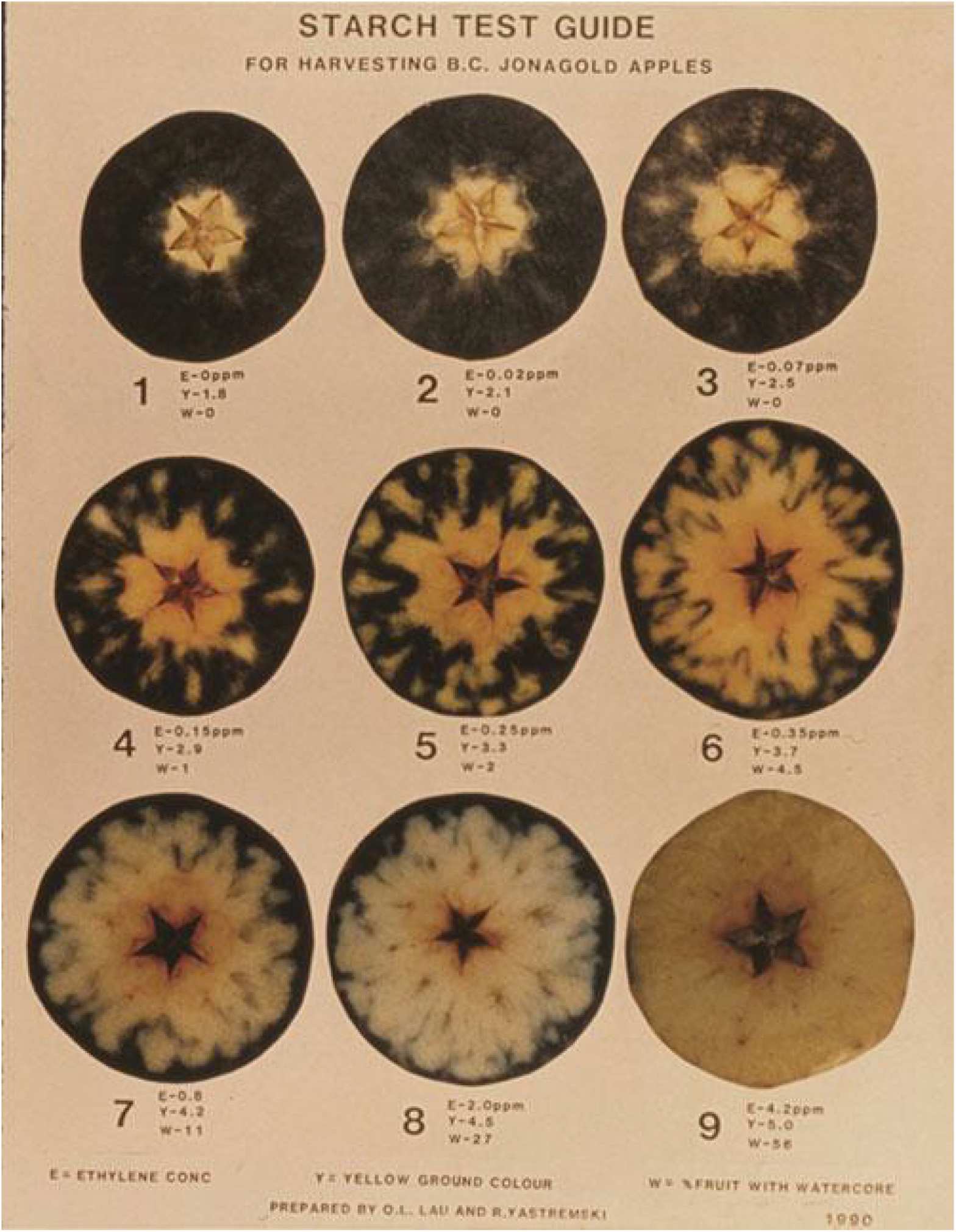
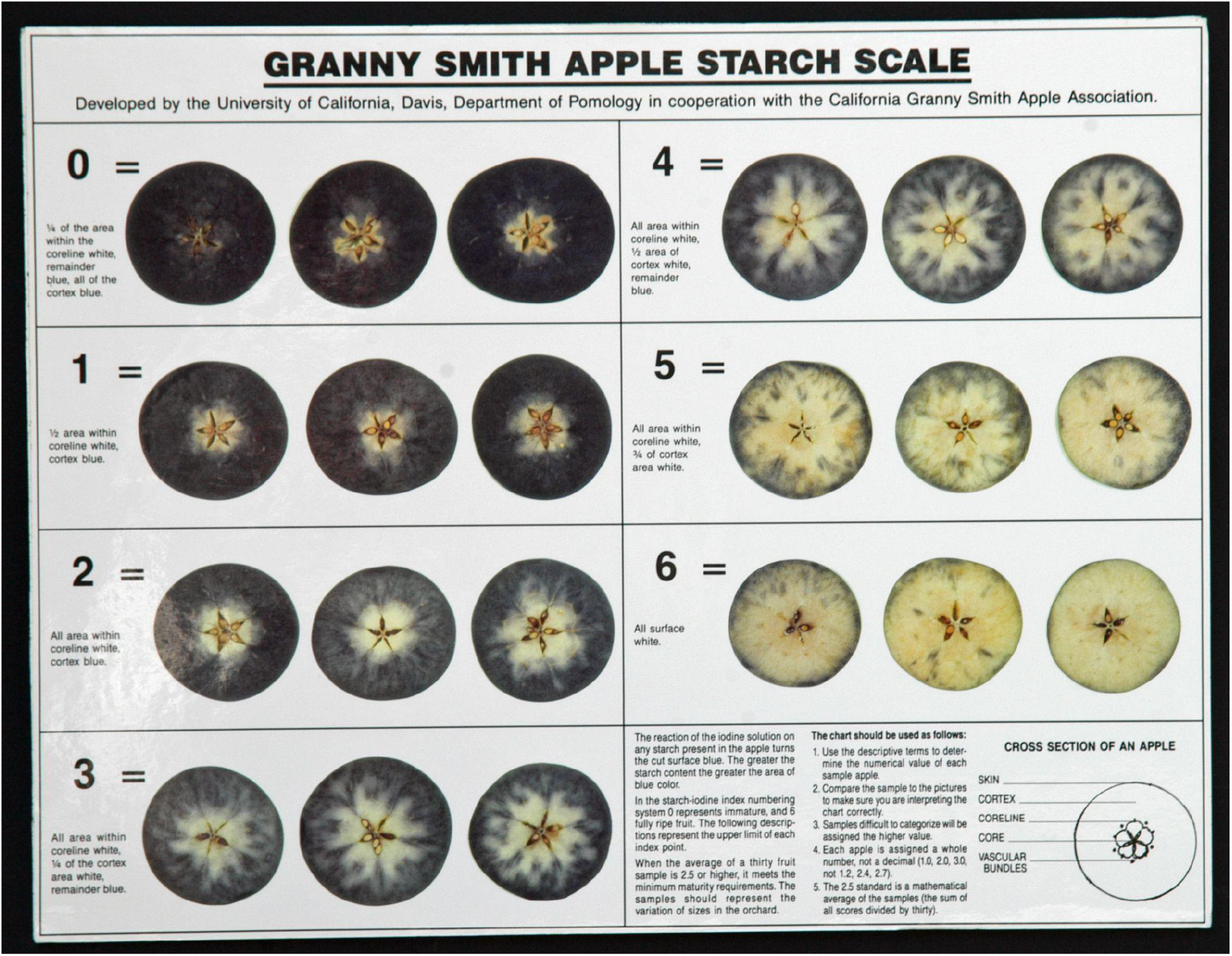
Starch pattern indices (SPI) commonly used for starch content assessment. From top to bottom, the SPIs are: The generic starch index card developed by Cornell University (as known as the Cornell Chart); The generic starch index card developed by ENZA Fruit; The starch index card designed for ‘Jonagold’ by O. L. Lau and R. Y. Yastremski; and the ‘Granny Smith’ starch scale developed by UC. Davis.

**Supplemental Figure 7.**
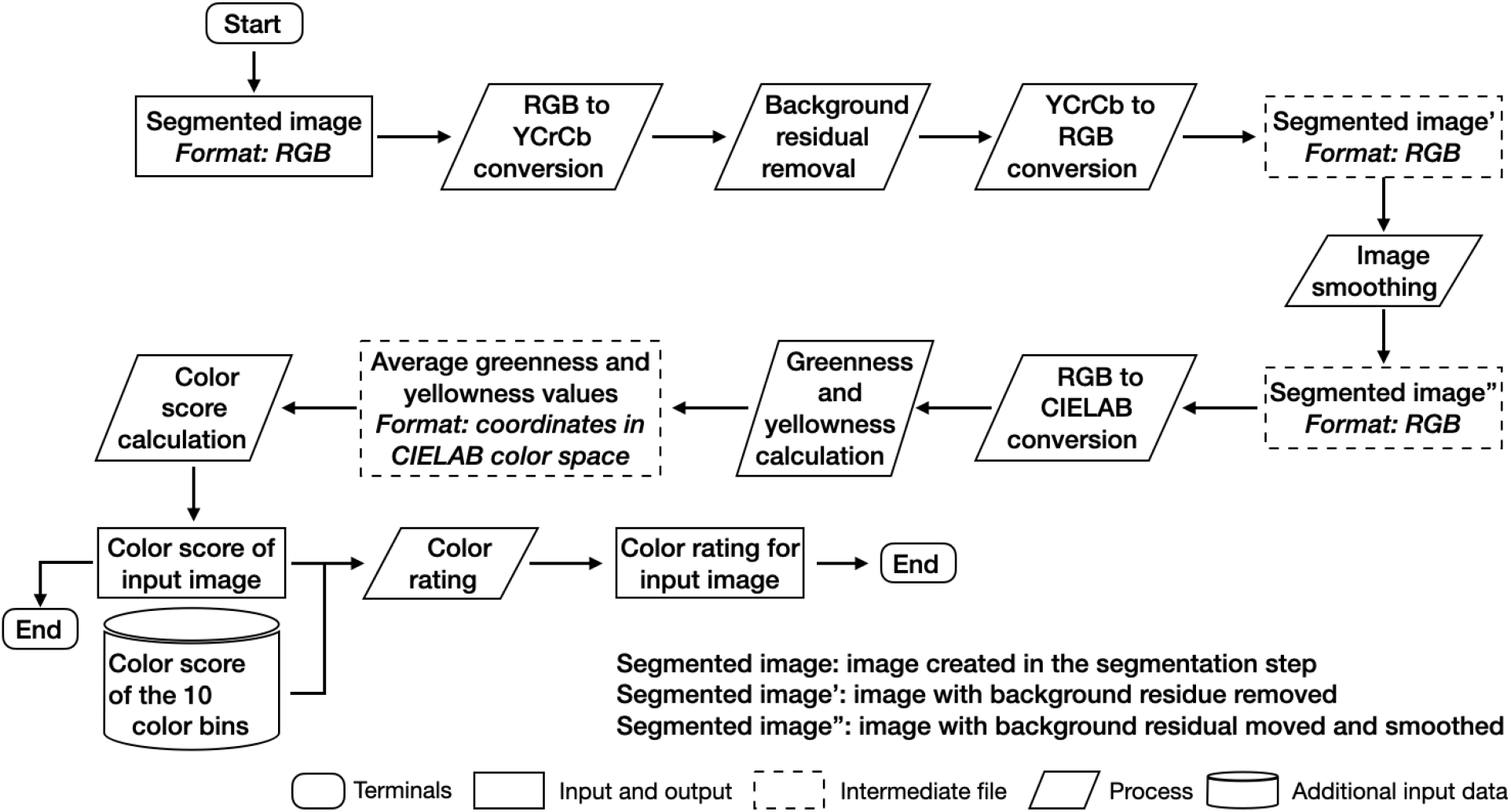
Pear background color rating workflow

**Supplemental Figure 8.**
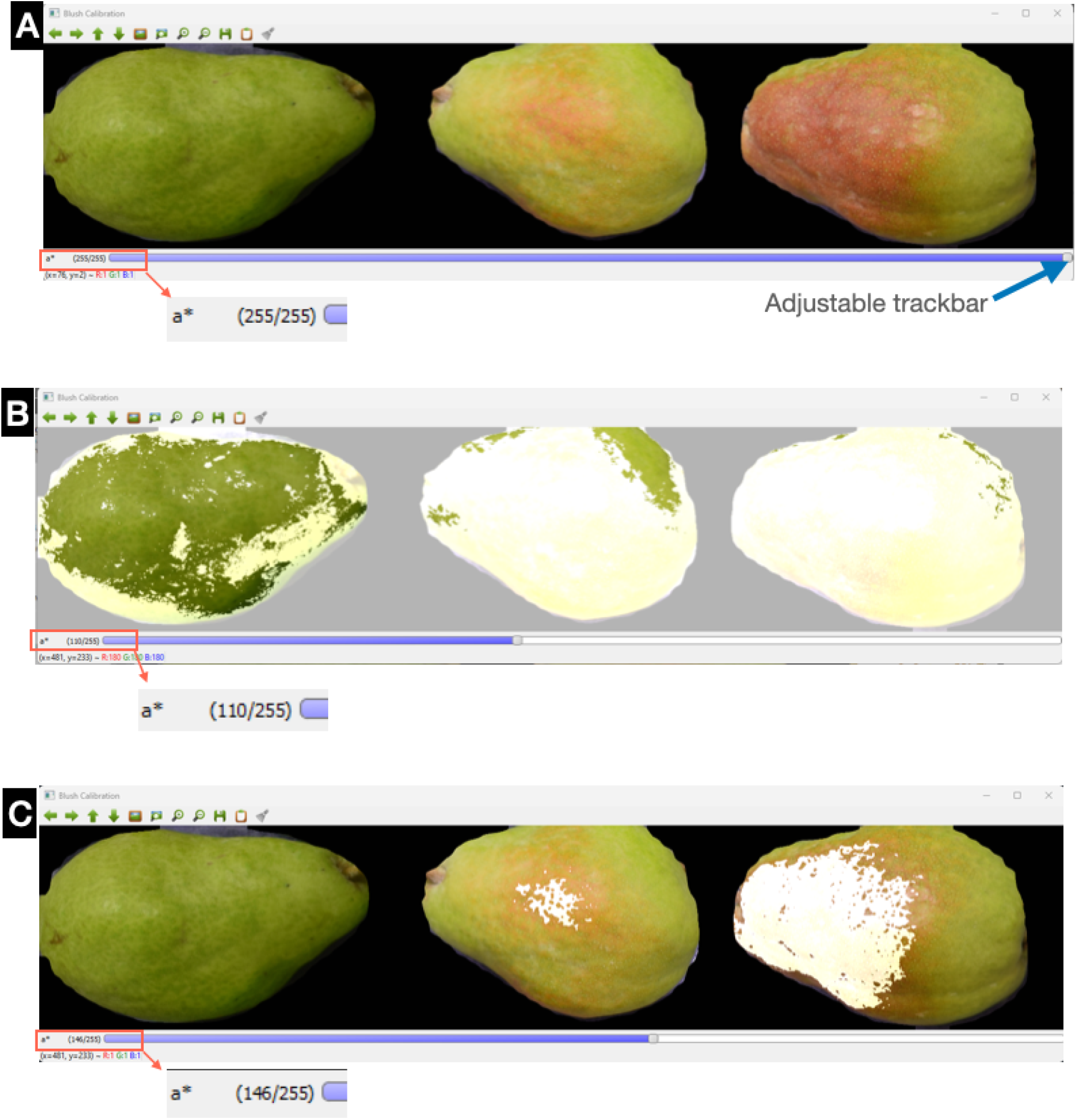
Example of the calibration step of the pear blush module visualization in an OpenCV window. **(A)** shows the three selected pear images representing no blush, light-colored blush, and intense-colored blush, from left to right. The trackbar, located on the bottom of the interface, can be used to adjust the threshold of the a* channel. The current a* channel reading and the max reading is shown to the left of the trackbar. **(B)** and **(C)** show the masks overlaying the pear images after adjusting the threshold using the trackbar. The mask in **(B)** covers only the blush region, but also green peel regions. The mask in **(C)** fails to cover all the blush regions. An ideal threshold would be between **(B)** and **(C)**.

**Supplemental Figure 9.**
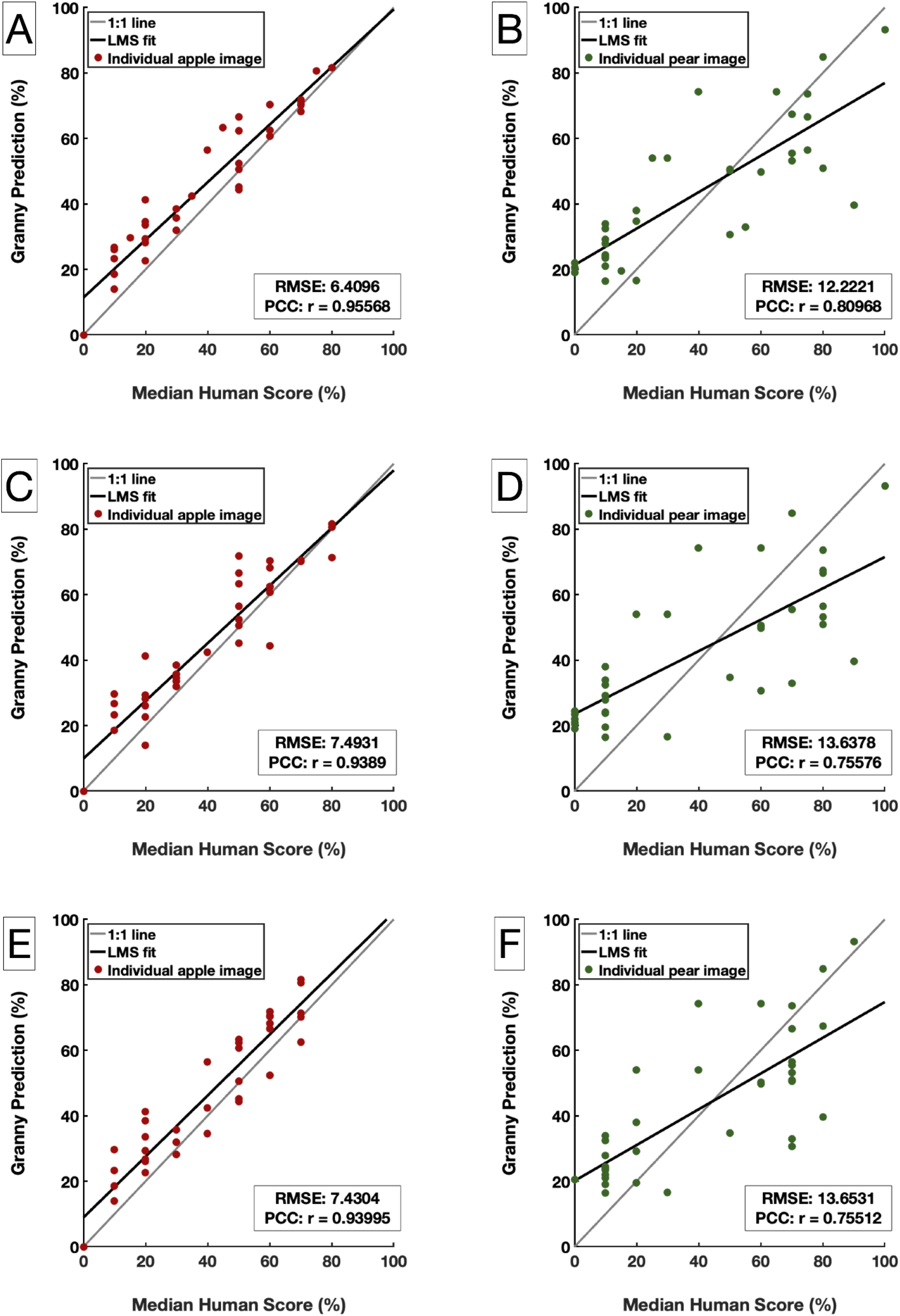
Starch content estimation from Granny and human participants with different levels of skills. (A), (C), (E) are score comparisons for apple cross-sections, while (B), (D), (F) are for pears. Participant skill levels are limited in (A) and (B), novice in (C) and (D), and experienced in (E) and (F). 36 apples from a variety of cultivars and 36 ‘Gem’ pears were evaluated. Numbers of participants in each skill group are: 18 participants have limited experience, 9 consider themselves novice technicians, and 9 are experts. LMS: Least Mean Square; RMSE: Root Mean Squared Error. PCC: Pearson Correlation Coefficient.

**Supplemental Figure 10.**
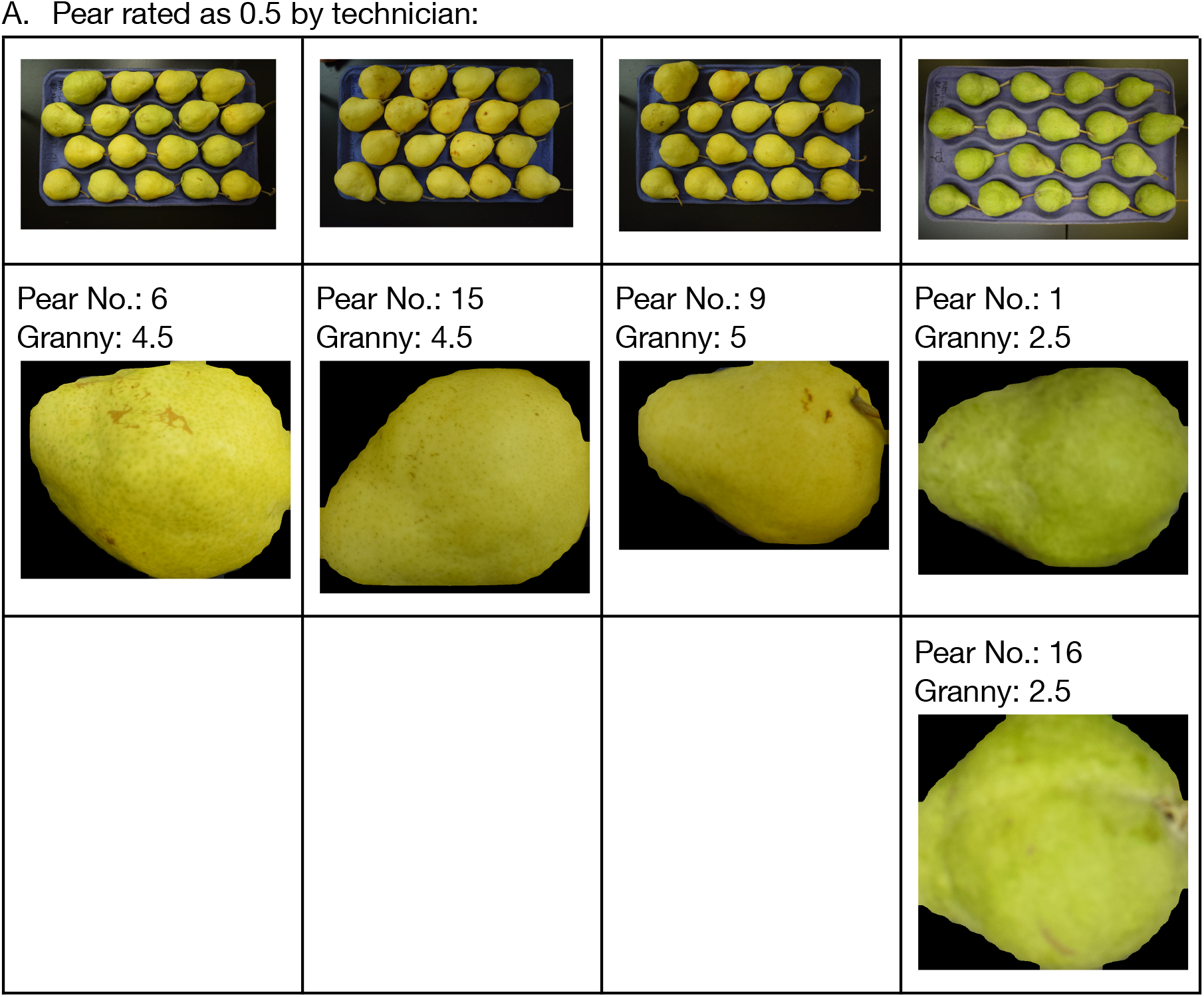
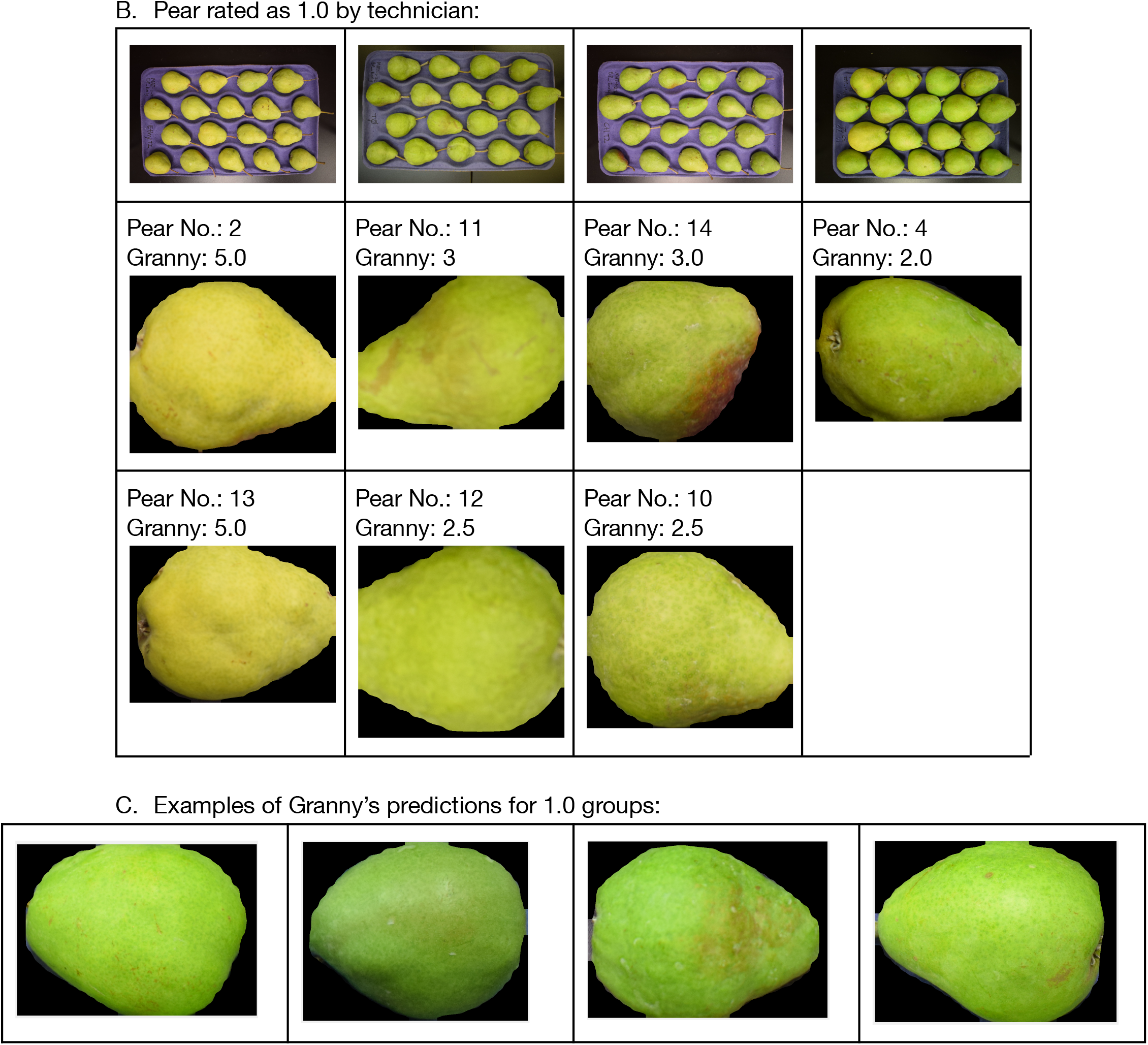
Examples of pears rated as 0.5 or 1 by technicians are shown in **(A)** and **(B);** examples of pears rated as 1 by Granny are shown in **(C)**. **(A)** and **(B)**: The top row contains images of the full try where the pears in question were extracted from. Pear are numbered from 1-18, starting from the top right corner. Second and third row are extracted pears from the corresponding tray, their location on the tray, and Granny’s color rating.

**Supplemental Figure 11.**
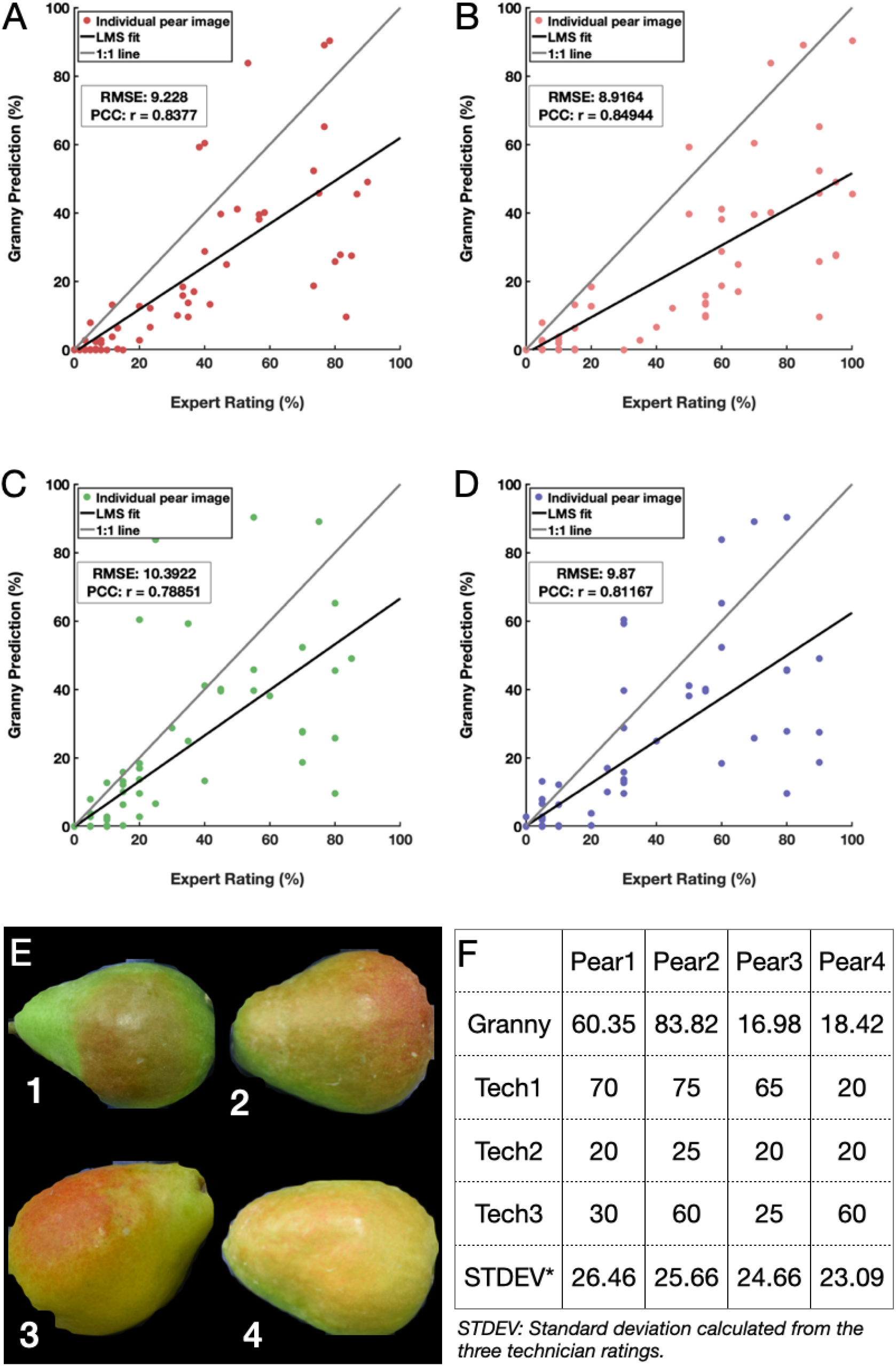
Pear blush rating from technicians compared to blush estimation from Granny. Granny predicted blush scores (y-axis) are compared to (A) the average rating from three technicians, (B) ratings from technician 1, (C) ratings from technician 2, and (D) ratings from technician 3. LMS: Least Mean Square; RMSE: Root Mean Squared Error. PCC: Pearson Correlation Coefficient. (E) and (F) are examples of the 4 pear images with the highest standard deviations from the three technician ratings.

## Supplemental Method - Starch Rating

The code of the ImageJ macro for starch rating is composed of six key regions: (1) a “fresh start” region where appropriate settings are applied and fields are reset for proper operation of the macro, (2) code which allows the user to choose input and output directories, (3) enabling of batch mode to fetch all images found in the input folder, (4) calculation of starch present in cross-sections, (5) calculation of total cross section area, and (6) saving data as a comma-separated file (CSV) to the output folder, where percent starch content is calculated per image.

When first initialized, appropriate settings are first applied to ImageJ ensuring accurate measurement of starch content. This includes (1) closing of any windows open in ImageJ, (2) resetting of the Region of Interest Manager (roiManager), (3) clearing data from the log file and subsequently closing the window, (4) setting the option “Black Background” to true, (5) enabling the display of file names for measurements, (6) setting the foreground color to white and background color to black, and (7) setting the width of drawn lines to one pixel. These settings must be changed for processes of the macro to run correctly.

Input and output directories are next designated by the user. The input folder should contain isolated cross-section images stained with iodine saved as a JPEG (.jpg), PNG (.png), or TIFF (.tif) format, with no limit on the number of images stored in the folder; all images of the input folder will be processed by the macro. The output folder is where threshold images of (a) starch area, (b) total cross-section area, and (c) the final comma-separated “Results.csv” file is saved. Threshold images of starch area and cross-section area are saved as a JPEG with the line “starch.jpg” or “totalarea.jpg” respectively added to the end of the original file name. After the input and output directories are chosen by the user, the macro will create a log in ImageJ with column titles which will be present in the final results file in comma-delimited format: data is added to this log following image analysis.

Batch mode is then enabled by the macro to fetch and process all images found in the input folder. Following this step all code is applied on a per-image basis; the script will repeat code until all images of the input folder have been processed. The macro will repeat code on subsequent images in the input folder until the last image is processed, where the log file is saved as “Results.csv” in the output folder.

Depending on the number of ROIs, the area of each region is identified by ImageJ as an alphabetical variable (C-L), which are later used by the macro to calculate starch content. The script is currently limited to ten identifiable ROIs, though could be modified to incorporate a higher number of regions. Based on our data set with apple and pear, ImageJ identifying more than six ROIs is uncommon.

Due to ImageJ’s region of interest (ROI) identification process, a thin line must be drawn by the macro to divide the image into two halves. If this step does not occur any images where starch runs the full circumference of the cross-section (similar in shape to a donut) the area calculated by ImageJ will be that of the inner non-iodine-stained area (Figure 6). Otherwise ImageJ identifies the inner region of such cross-sections as the primary region-of-interest (ROI), instead of the actual iodine-stained region. With the image divided into two halves, the iodine-stained area of the image is correctly identified since no single ROI can run the full circumference of the cross-section.

**Figure S1.**
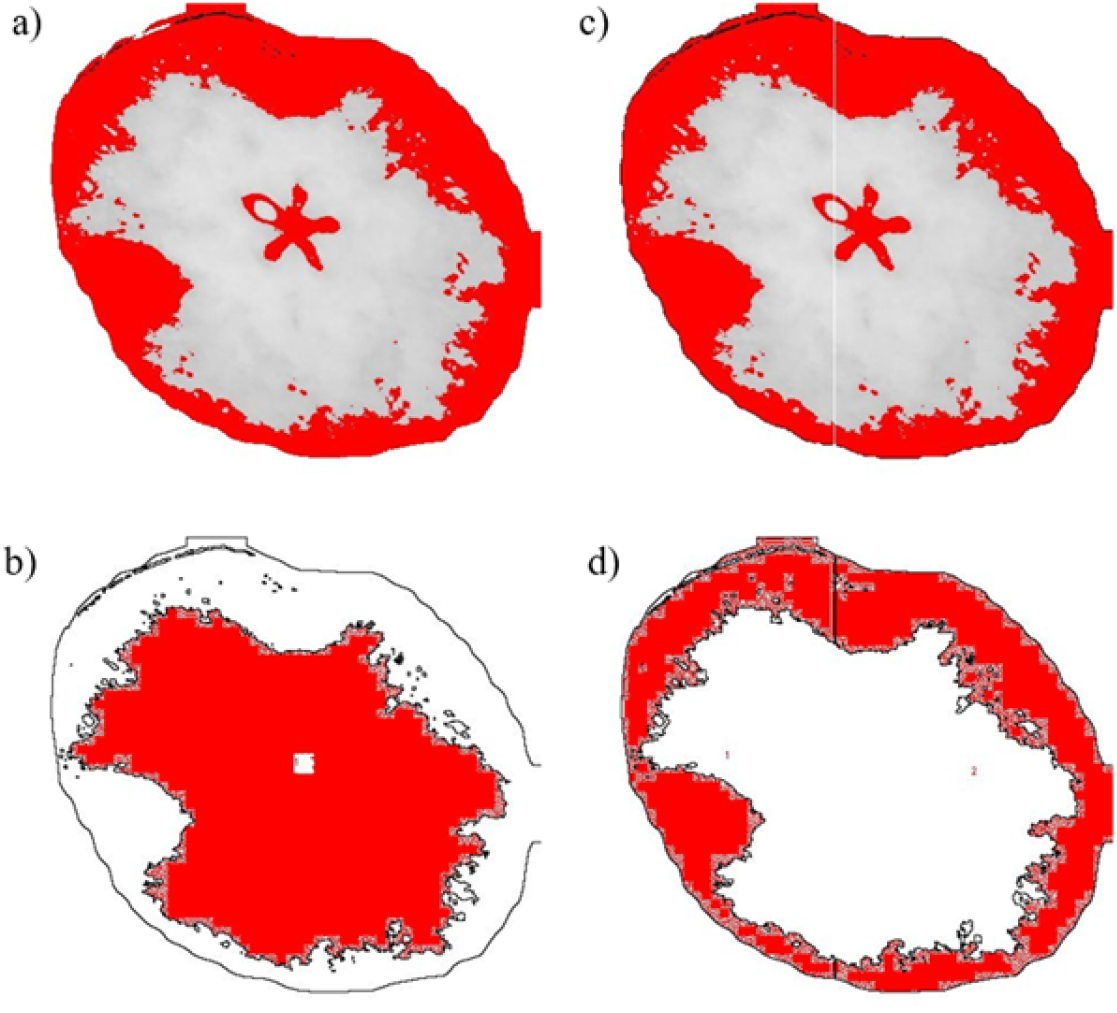
Starch analysis images thresholded with (a) and without (c) a division line, and identified ROIs of the above threshold images with (b) and without (d) a division line. Without a division line one incorrect ROI was identified (b), while with a division line (d) two correct ROIs were identified. Iodine-stained regions were correctly identified in thresholds with a division line (c, d). A starch threshold image is saved to the output folder as a JPEG with “starch.jpg” appended to the image’s original file name. The macro will then pause for 100 milliseconds permitting file saving on slower systems. After another 100 millisecond pause, a second threshold is then applied to the image to calculate the total area of the cross-section. Using the “Analyze Particles” function, the total area of the cross-section threshold is calculated, and the threshold is saved to the output folder with “total_area.jpg” appended to the image’s original file name. The total area of the cross section is saved by ImageJ as variable “B”. The code then assesses the number of ROIs, calculates the sum ROI area, and divides the sum ROI area by the total cross-section area to calculate percent starch area of the cross-section, saving this data to the macro’s log file. Using percent starch content best-fit polynomial lines of established rating scales, the starch rating for each image is also calculated and rounded to the nearest integer by the macro. Established rating scales calculated include: the Cornell Chart (Blanpied and Silsby 1992), Jonagold (REF), Purdue (REF), and University of California - Davis’ ‘Granny Smith’ starch rating scales (Mitcham et al 1996). These values are found in the designated output folder “Results.csv” file.

**Figure S2.**
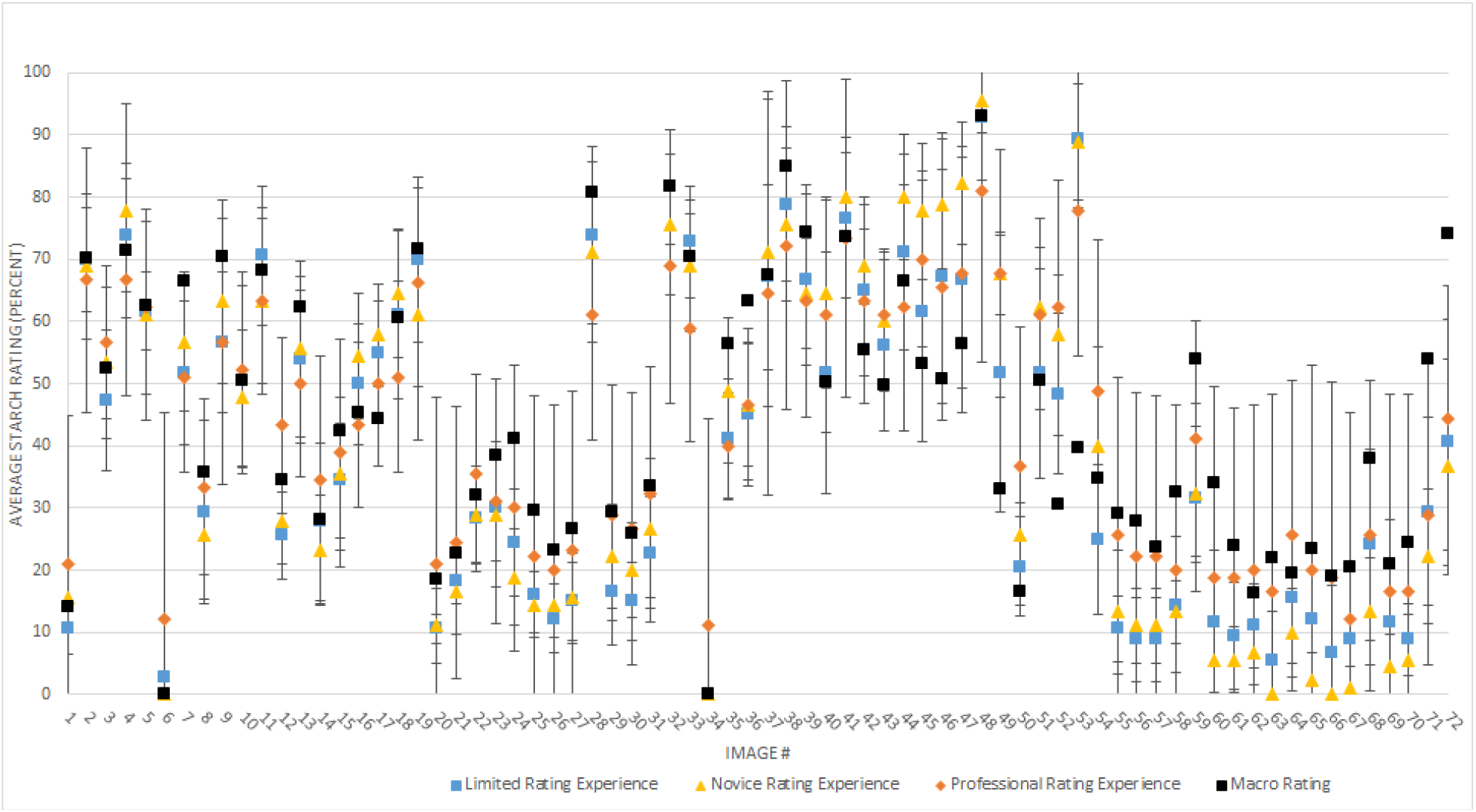
Average percent starch rating of the limited rating experience rating (blue, N = 18), novice rating experience (yellow, N = 9), professional rating experience (orange, N = 9), and macro starch rating output (black). Images 1-36 iodine-stained ‘Granny Smith cross-sections. Images 37-72 iodine-stained ‘Gem’ pear cross-sections. (https://drive.google.com/drive/u/0/folders/1PD_WQrnZfrZYaZVfgNmpeVp0a_Bdx5o5)

**Figure S3.**
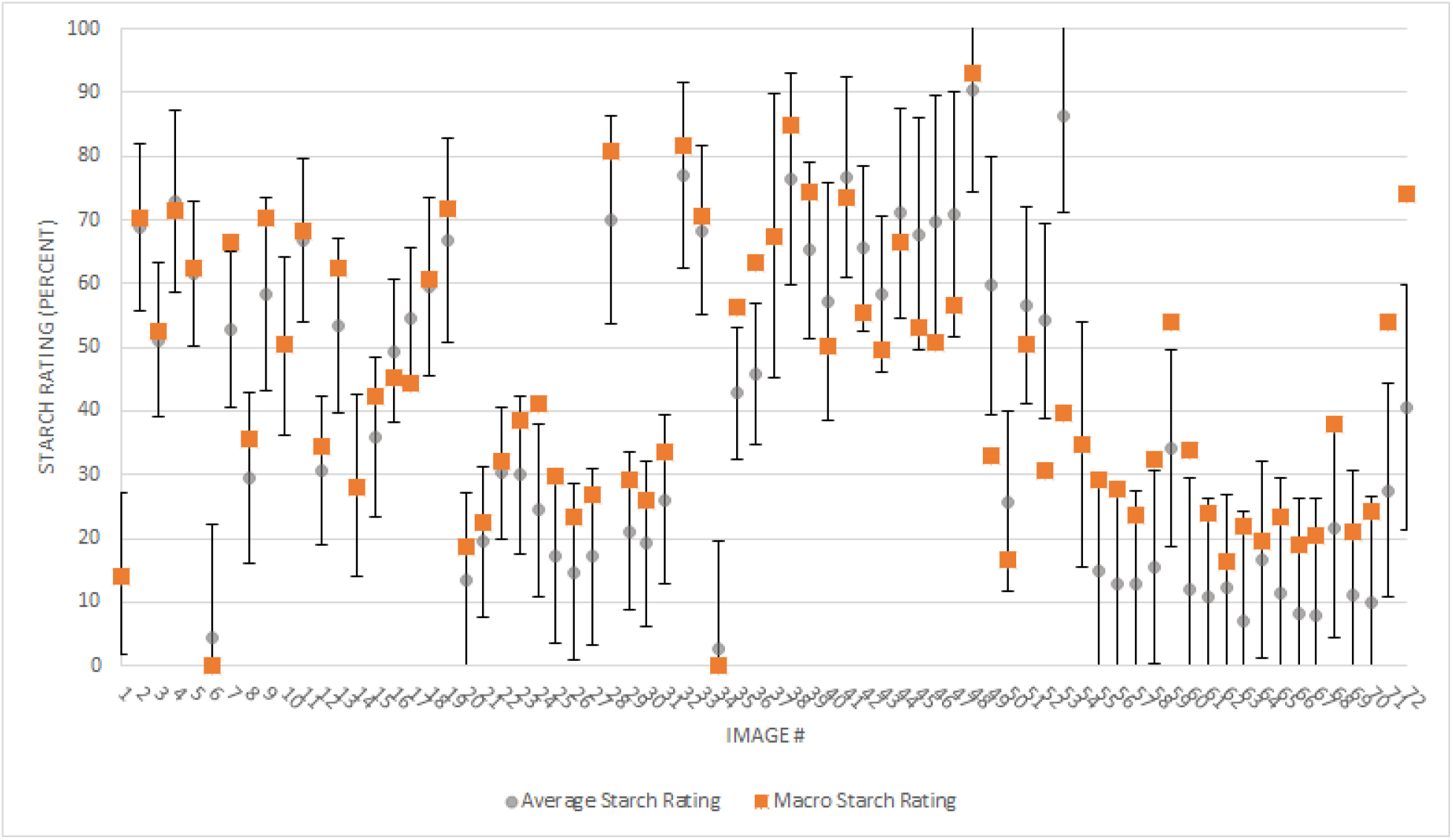
Average percent starch rating of all participants of the visual starch rating assessment (black, N = 36) compared to macro starch rating output (orange). Images 1-36 iodine-stained ‘Granny Smith’ apple cross-sections. Images 37-72 iodine-stained ‘Gem’ pear cross-sections. (https://drive.google.com/drive/u/0/folders/1PD_WQrnZfrZYaZVfgNmpeVp0a_Bdx5o5)

